# Population-scale measurement of human centromere array length variation from short-read sequencing

**DOI:** 10.64898/2026.11.25.769230

**Authors:** Yikai Dong, Joshua Weinstock

## Abstract

Alpha-satellite higher-order repeat (HOR) arrays at human centromeres vary several-fold in length across individuals and have been linked to kinetochore function and chromosome missegregation. However, their repetitive megabase-scale architecture has largely restricted length measurement to a few hundred long-read derived genome assemblies, leaving short-read cohorts uncharacterized. Here we introduce cenlen, an alignment-free method that estimates active centromeric array lengths across all 24 human chromosomes directly from short- read or long-read CRAM files. To estimate array lengths, we first identify *k*-mers that are common within a query chromosome’s centromere array but rare across other chromosomes. for acrocentric and other repeat- sharing families lacking private k-mers, cenlen instead models shared k-mer mixtures against assembled pangenome reference profiles using non-negative linear deconvolution. Across 61 genomes that were not used for training, cenlen estimates agreed closely with assembled array lengths (median relative error 2.4%), outperforming existing short-read estimators. We further optimized sequence retrieval and targeted k-mer counting, reducing CPU time 313-fold relative to existing methods. We applied cenlen to 2,557 individuals across genetic ancestries from the 1000 Genomes Project. Most variation in array length occurred within populations, but arrays on chr21, chr7 and chr16 differed between superpopulations more than the average common SNV. finally, we found that array lengths were associated with molecular phenotypes, including splicing of *MED14* in chromosome X. Overall, cenlen makes it possible to test associations between centromere array length variation and human traits in large sequencing cohorts.

## Introduction

Human centromeres are defined by megabase-scale arrays of alpha-satellite higher-order repeats (HORs) [1]. The active array of each chromosome varies several-fold in length between homologues and across populations [2–4], and array size has been linked to kinetochore assembly [5] and chromosome missegregation [6, 7]. Recent work has now resolved telomere-to-telomere assemblies, including centromere array sequence in hundreds of individuals [1, 4, 8]. Although an exceptional achievement, these sample sizes remain relatively small within the scope of human genetics association studies. If lessons from other genetic association studies are to follow, then we may need thousands of centromere sequences if we are to resolve connections between centromeric variant and molecular, cellular, and organismal phenotypes.

In recent years, population biobanks have sequenced cohorts primarily with short reads [9, 10] across enormous numbers of individuals (400K∼500K). We recently showed that centromeric structural variants can be statistically imputed into these short-read genomes and thus can be used in large-scale association studies [11]; however, this work does not clarify the role of centromere array lengths.

Although short reads collapse repetitive arrays during alignment, their sequencing depth remains proportional to underlying array length, enabling abundance-based length estimation from array-specific k-mers [2]. This principle was established on individual sex chromosomes [1, 2, 12] and recently extended to 17 autosomes by HOROSCOPE [13]. Profiling every chromosome from short reads poses two further challenges. first, the active arrays of chr13 and chr21, of chr14 and chr22, and of chr1, chr5, chr16 and chr19 share near-identical HOR units across chromosomes through concerted evolution [14, 15]; these arrays contain few chromosome-unique k-mers, and the acrocentric arrays of chr13, chr14, chr21 and chr22 lie outside the scope of existing estimators [13]. Second, existing workflows decompress entire alignment files to count k-mers genome-wide, which requires millions of CPU hours and large-volume data transfers to profile biobank cohorts, thus making the application of existing approaches cost prohibitive.

To address these challenges, we developed cenlen, an algorithm that estimates active centromere lengths across all chromosomes directly from short-read CRAM files (fig. 1). cenlen uses the CRAM index to retrieve containers overlapping targeted regions, including centromeric arrays and single-copy normalization regions. It then decodes read flags, reference identifiers and bases to count k-mers. To resolve arrays that share HOR units, cenlen models normalized k-mer abundances as non-negative linear combinations of assembled pangenome array profiles and reports diploid lengths with empirically calibrated prediction intervals.

**Figure 1:**
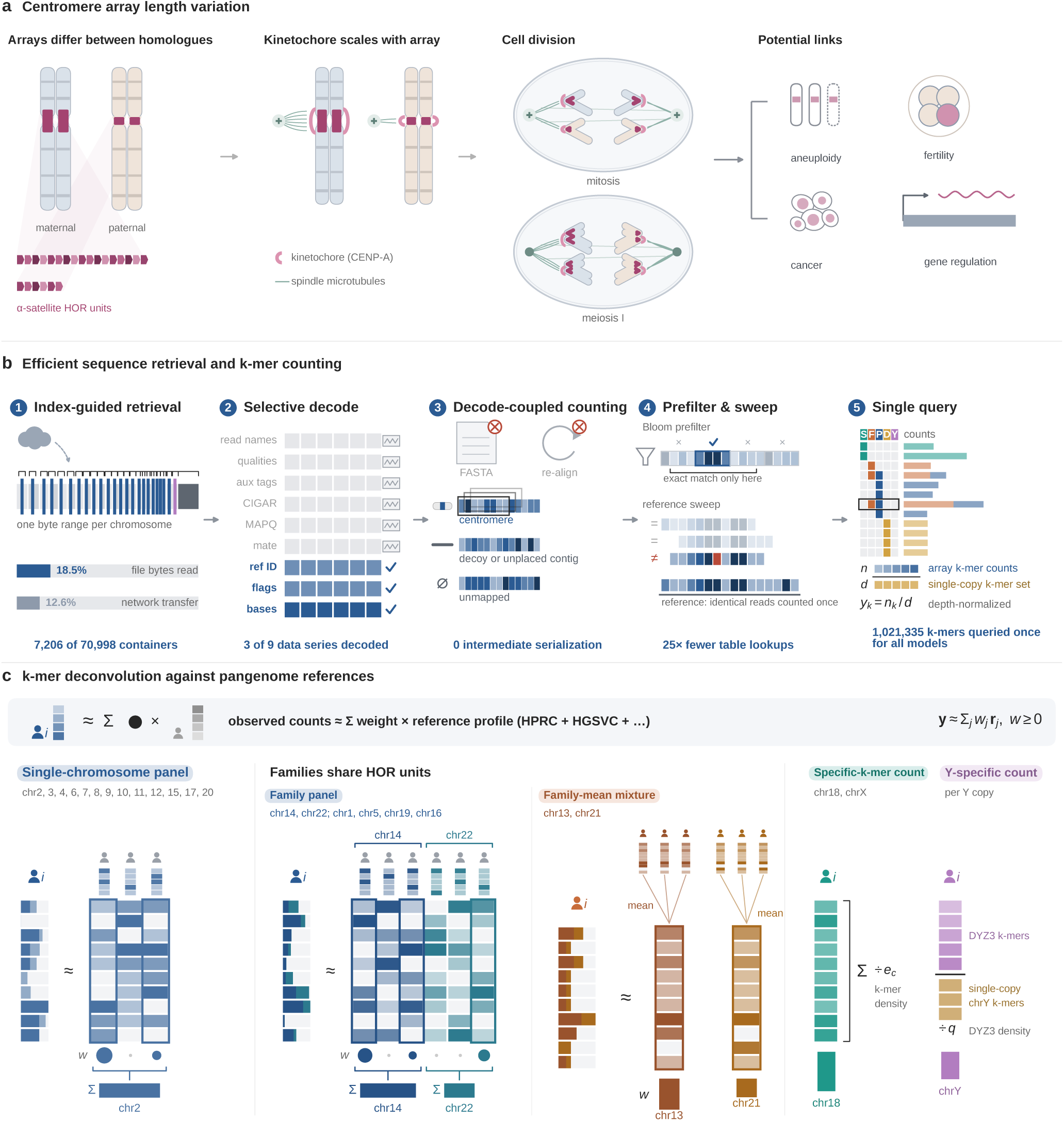
Centromere array length, targeted CRAM decoding and pangenome-referenced length estimation. (a) Active alpha-satellite HOR arrays differ in length between homologues and individuals; array length is linked to kinetochore size and missegregation, and potentially to aneuploidy, fertility, cancer and gene regulation. (b) cenlen reads only the CRAM containers that overlap 3,432 target intervals, decodes three of nine data series, counts mapped, unplaced and unmapped reads in one decode-coupled pass, screens reads with a 16-base Bloom prefilter and counts reference-identical reads once per slice, and queries each canonical 31-mer once against 1,021,335 indexed k-mers; bit columns mark chromosome-specific sets (S), family sets (f), panel groups (P), single-copy depth k-mers (D) and chrY sets (Y), and counts *n* divided by the per-copy depth *d* give abundances *y_k_* = *n_k_*/*d*. (c) Each model fits the abundances of genome *i* as a non-negative weighted sum of assembled reference profiles: the single-chromosome panel against the individual arrays of one chromosome (13 chromosomes) and the family panel against those of a shared-HOR family (chr14 and chr22; chr1, chr5 and chr19 with chr16); the family-mean mixture of member mean profiles (chr13, chr21); the specific-k-mer count, a chromosome-specific k-mer sum divided by its density *e_c_* (chr18, chrX); and the Y-specific count, DYZ3 counts per single-copy chrY k-mer divided by the DYZ3 density *q* (chrY). Circles, fitted weights *w* (area proportional to weight; grey, zero). Colours mark the four models throughout.

Across 61 validation genomes, 427 pedigree trios and 2,557 unrelated individuals of the 1000 Genomes Project, cenlen achieves assembly-concordant accuracy. Applied to global diversity and functional genomics cohorts, it reveals population-stratified array lengths, including a longer chr21 array in the African superpopulation, and an association of X-array length with *MED14* splicing, establishing centromere array length as a tractable and important source of genetic variation in large-scale genetic association studies.

## Results

Centromere arrays are highly repetitive and thus difficult to study through standard alignment based approaches. An alternative to alignment based approaches is to use k-mers, which are continuous DNA substrings. We reasoned that it may be possible to compute k-mers that are specific to, or at least enriched for, centromere repeat arrays. Recently, the Human Pangenome Reference Consortium (HPRC) and Human Genome Structural Variation Consortium (HGSVC) have used a combination of Nanopore, Pacbio and other long-read technologies along with algorithmic innovations to assemble a few hundreds genomes. Many of these assemblies include contigs that even span centromere arrays. We reasoned that these centromere-spanning assemblies could serve as a reference for identifying specific k-mers that tag centromere arrays lengths.

To assess this hypothesis, we searched for canonical 31-mers that were useful for distinguishing centromere arrays in the HPRC and HGSVC assemblies. Specifically, we searched for 31-mers that were specific to either single- chromosome arrays or specific families of chromosomes with highly similar arrays (e.g., chr13 and chr21, chr14 and chr22). We then constructed a statistical model that used these array fingerprint 31-mers as predictors for the array lengths that were derived from the genome assemblies. Array lengths from homologous chromosomes were then summed. To adjust for sequencing depth, we curated a distinct set of k-mers that are useful for estimating depth at a prespecified set of loci (Methods). We refer to this approach as *cenlen*.

cenlen estimates the active array length of every chromosome from 18.5% of a CRAM file’s bytes (fig. 1). It decodes only the containers that overlap 3,432 target intervals and counts each canonical 31-mer once against an indexed table of 1,021,335 31-mers. We then normalized counts by the depth of autosomal k-mers from the same intervals (Methods).

We then trained models to estimate array lengths from the normalized counts using the HPRC/HGSVC genome assemblies to define the ground truth. We developed distinct models for the pairs of chromosomes with highly similar arrays (i.e., chr13-chr21, along with sex-chromosome specific approaches). We developed a reference-panel approach, where we fit a non-negative weighted-least squares model to a reference panel of assembled haplotype array profiles (fig. 1c; Methods).

### cenlen estimates array lengths across all chromosomes, including those with shared repeats

To validate the *cenlen* estimates of array lengths, we used a distinct set of genome assemblies to test its performance. We computed the chromosome 31-mer fingerprints and used them to generate array lengths from our statistical model. In the 61 validation genomes with completely assembled arrays, Pearson *r* ranged from 0.88 to 0.999 and the median relative error (MedRE) across targets was 2.4% (2.7% pooled over all genome–array pairs), with 23 targets at or below 5% (fig. 2a; Supplementary fig. S1; Supplementary Table S1). We also asked whether our uncertainty estimates were well calibrated by quantifying empirical coverage. We observed that the prediction intervals were calibrated, with 95% intervals including 94.3% of the 566 assembled lengths (Supplementary fig. S2a), with a median width of 19.9% of the estimate. The widest intervals belong to chr13 and chr21 (67.2% and 77.0%), whose arrays share 96–97% of their k-mer copies (k-mer occurrences counted with copy number; family total in fig. 2f).

**Figure 2:**
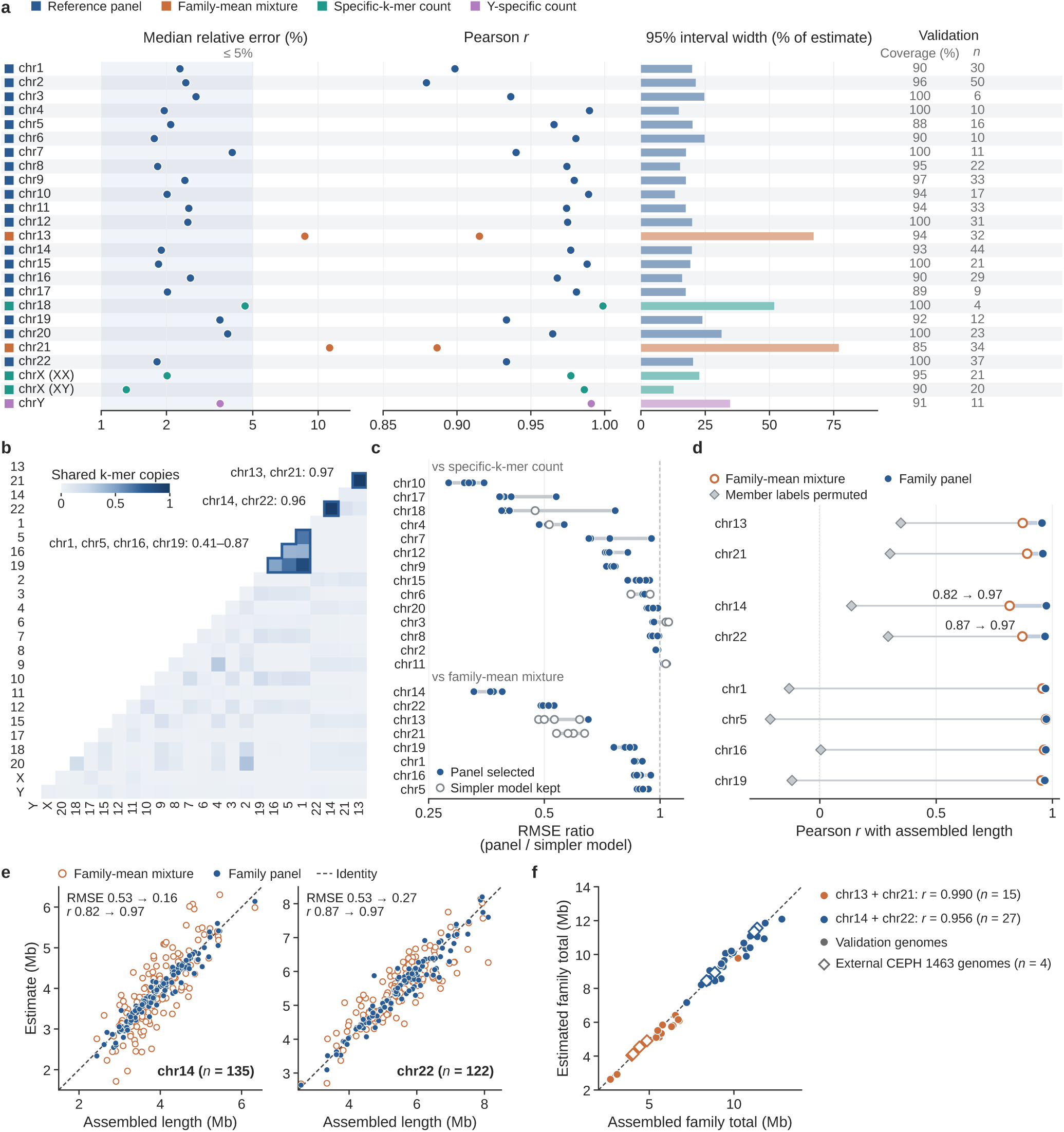
Accuracy across all 25 targets and deconvolution of shared-HOR families. (a) Per-target accuracy on the 61 validation genomes (566 genome–array pairs; nested cross-validation in Supplementary Table S2); squares, model class: median relative error (shaded, ≤ 5%), Pearson *r* and median 95% prediction-interval width (% of the estimate; bars, validation; median 19.9% overall and 19.8% on panel targets, range 13.2–31.3%); right, validation 95% coverage and sample sizes. (b) Shared k-mer copies between active arrays (larger directed share) across training assemblies; framed, the three repeat-sharing families, the only pairs sharing at least 0.40. (c) RMSE of the reference panel relative to the simpler model (specific-k-mer count or family-mean mixture) per outer fold of nested cross-validation; filled, folds where the automated rule selected the panel. (d) Correlation with assembled length for the eight family members under the family-mean mixture, the family panel and family panels with member labels permuted, in family-grouped cross-validation on short reads. (e) chr14 and chr22 in the same cross-validation, with RMSE (Mb) and *r* of both models; dashed line, identity. (f) family totals in the validation genomes (filled) and the CEPH 1463 genomes (open diamonds).

We performed a second validation using six CEPH 1463 genomes that were not included in cenlen’s training or initial validation sets. Our estimates on these genomes had a median relative error of 1.2% with 82 of 83 lengths inside the 95% intervals (Supplementary fig. S5a, Supplementary Table S2; Supplementary fig. S5b).

Because the number of informative k-mers per unit of array length can vary across haplotypes, a single average k-mer density may not accurately represent every array. We therefore tested whether a panel of assembled arrays improved length estimation. On chr7, chr10, and chr22, the reference panel lowered cross-validated RMSE from 0.34, 0.45, and 0.53 Mb under simpler models to 0.27, 0.14, and 0.27 Mb (fig. 2c; Supplementary Section S5).

Because some groups of chromosomes have highly similar arrays, we then asked which of our modeling approaches had the strongest performance. for chr14 and chr22, jointly modeling the two arrays with a reference panel increased the correlation with assembled length from 0.82 to 0.97 and from 0.87 to 0.97, respectively, compared with the family-mean mixture. for chr13 and chr21, the individual arrays have the largest errors, and their sum, like that of chr14 and chr22, matched assembled lengths closely (fig. 2a,f).

### Pedigree transmission, comparisons with existing methods and down-sampling confirm fidelity

As an orthogonal validation, we asked whether our array estimates conformed to Mendelian inheritance expectations within sequenced trios. Across 427 parent–offspring trios of the high-coverage 1000 Genomes cohort, the transmission slope *R* of the 22 autosomal arrays had a median of 0.955 (range 0.854–1.131), against upper 95% limits of at most 0.30 with children permuted within populations (fig. 3a; Supplementary Table S1). The sex-chromosome arrays followed their expected inheritance on the per-copy scale: 1.09 for a son’s X on the mother’s (expected 1), 0.52 for a daughter’s X on the father’s (expected 0.5) and 0.991 for the Y array across 308 father–son pairs. Taken together, these results confirm that our array length estimates conform to Mendelian inheritance expectations in both autosomes and sex chromosomes.

**Figure 3:**
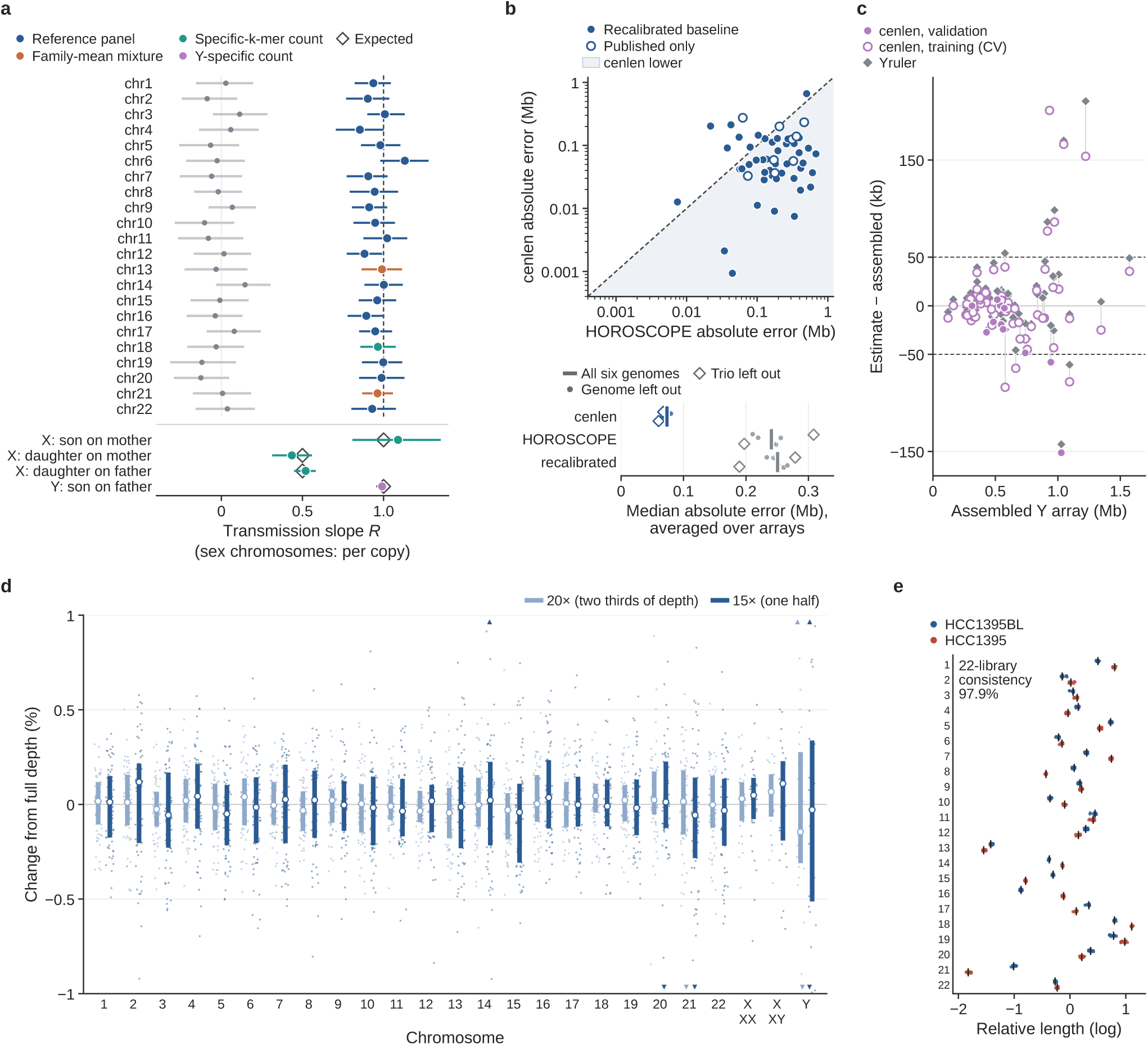
Transmission in families, comparison with existing methods and robustness to sequencing depth and centre. (a) Mendelian transmission slope *R* of the 22 autosomal arrays across 427 parent–offspring trios (dots coloured by model; 95% bootstrap intervals), beside *R* with children permuted within populations (grey); dashed line, *R* = 1. Below, sex-chromosome arrays on the per-copy scale; diamonds, expected slopes. (b) Six CEPH 1463 genomes against Platinum Pedigree assemblies: absolute error of cenlen against HOROSCOPE (56 genome–array pairs; shaded, cenlen lower; filled, recalibrated baseline). Strip, array-averaged median absolute error: all six genomes (bars), each genome left out (dots), each trio left out (diamonds). (c) Y array in 70 XY genomes: estimate minus assembled length for cenlen (filled, validation; open, cross-validated training) and Yruler, joined per genome; dashed lines, ±50 kb. (d) Change of each estimate after down-sampling the 61 validation libraries to 20× and 15× (bars, interquartile range; dots, medians; triangles, the 8 of 2,860 changes beyond ±1%). (e) Same DNA across sequencing centres: relative length of each autosomal array in 22 PCR-free libraries of HCC1395BL (blue) and HCC1395 (red), the lymphoblastoid and tumour lines of one donor; points, libraries; bars, medians (Supplementary Section S12).

We next benchmarked accuracy against existing short-read estimators using independent reference assemblies (Table 1). In the six CEPH 1463 genomes with Platinum Pedigree assemblies [16], cenlen reduced median absolute error more than threefold relative to HOROSCOPE [13] (fig. 3b; Supplementary Table S6). Notably, cenlen achieved the lower error in 47 of the 56 assemblies.

**Table 1:** cenlen, existing short-read estimators and long-read assemblies. MedRE, median absolute relative error; MedAE, array-averaged median absolute error; MedAPE, median absolute percentage error; CEPH 1463, six genomes with Platinum Pedigree assemblies; –, not applicable. CPU times were measured on the same compute node for each comparison (Supplementary Table S12).

| **Capability** | **cenlen** | **HOROSCOPE** | **Yruler** | **Long-read assembly** |
|---|---|---|---|---|
| Chromosomes | All 24 (25 targets)<sup>a</sup> | 17 autosomes | chrY | All 24 |
| Shared-HOR families | Deconvolved by panel and mixture models<sup>b</sup> | chr13, chr14, chr21, chr22 not estimated | – | Resolved by assembly |
| Prediction intervals | Calibrated 90% and 95%<sup>c</sup> | – | – | – |
| chrY readout | DYZ3 length per Y copy; Y dosage<sup>d</sup> | – | DYZ3 length | DYZ3 length |
| Accuracy | MedRE 2.4% (61 genomes); MedAE 0.073 Mb<sup>e</sup>; chrY MedAPE 2.8%<sup>f</sup> | MedAE 0.241 Mb published, 0.251 Mb recalibrated<sup>e</sup> | chrY MedAPE 2.5%<sup>f</sup> | Reference |
| CPU time, 30× genome | 42 s (25 targets); 35 s (17 autosomes); 6.2 s (chrY); 0.65 GB memory | 11,060 s (17 autosomes) | 49.6 s (region); 939 s (whole file) | Long-read sequencing and assembly |
| Data read | 18.5% of file bytes locally; 12.6% over HTTP (DRAGEN) | Whole file | Region or whole file | Long reads |

<sup>a</sup>22 autosomes, chrX (XX), chrX (XY) and chrY. <sup>b</sup>chr13 and chr21; chr14 and chr22; chr1, chr5 and chr19 with chr16. <sup>c</sup>Coverage 89.4% and 94.3% in 61 validation genomes; 91.1% and 96.2% in nested cross-validation. <sup>d</sup> *R*<sub>*Y*</sub> = *u*<sub>*Y*</sub>/*d* reads out Y dosage (somatic loss of Y) separately from array length; XX maximum 0.0078 and XY minimum 0.396 across 3,201 classified genomes. <sup>e</sup>CEPH 1463, 47 genome–array pairs with a recalibrated HOROSCOPE baseline; 0.085 against 0.237 Mb across all 56 pairs. <sup>f</sup>70 XY genomes; RMSE 50 kb (cenlen) and 51 kb (Yruler).

On chrY, cenlen had similar performance to Yruler [12] while also measuring Y dosage (fig. 3c; Supplementary fig. S7; Supplementary Table S7). Across all 70 XY genomes with assembled arrays, the two methods were statistically indistinguishable (median absolute percentage error 2.8% and 2.5%; RMSE 50 and 51 kb; paired Wilcoxon *p* = 0.79). Lastly, we showed that cenlen estimates were highly consistent across sequencing depths, platforms, and library preparations (fig. 3d,e; Supplementary Table S8; Supplementary Section S12). Down-sampling validation libraries to 15×–20× barely altered accuracy (pooled MedRE remained 2.7%; 95% interval coverage remained 94%). We evaluated reproducibility across 103 SEQC2 libraries prepared using several protocols. Among the PCR-free libraries, relative array-length estimates were highly consistent across six sequencing centers and three Illumina platforms, with a median reliability of 0.99 (fig. 3e).

### Targeted decoding and decode-coupled counting significantly reduce CPU time and scale to large cohorts

Estimating array lengths in hundreds of thousands of genomes requires an efficient way to retrieve and count the relevant reads. We therefore designed cenlen to retrieve targeted CRAM containers and decode the fields needed for k-mer counting. Restricting retrieval to targeted containers and decoding only three of nine CRAM data series reduced CPU time 11-fold relative to whole-file streaming through conventional tools (samtools), and decode- coupled counting reduced CPU time a further 4-fold, with identical k-mer counts (Supplementary Table S13). Read names, alignment tags and base qualities occupy 80.7% of a 30× CRAM file and sequence bases 7.5% (fig. 4a); by selecting containers through the index and decoding only read flags, reference identifiers and bases, cenlen reads 18.5% of file bytes locally (17.1–20.7% across seven 1000 Genomes files) and 12.6% over HTTP on *All of Us* DRAGEN alignments (Supplementary Sections S2 and S8).

**Figure 4:**
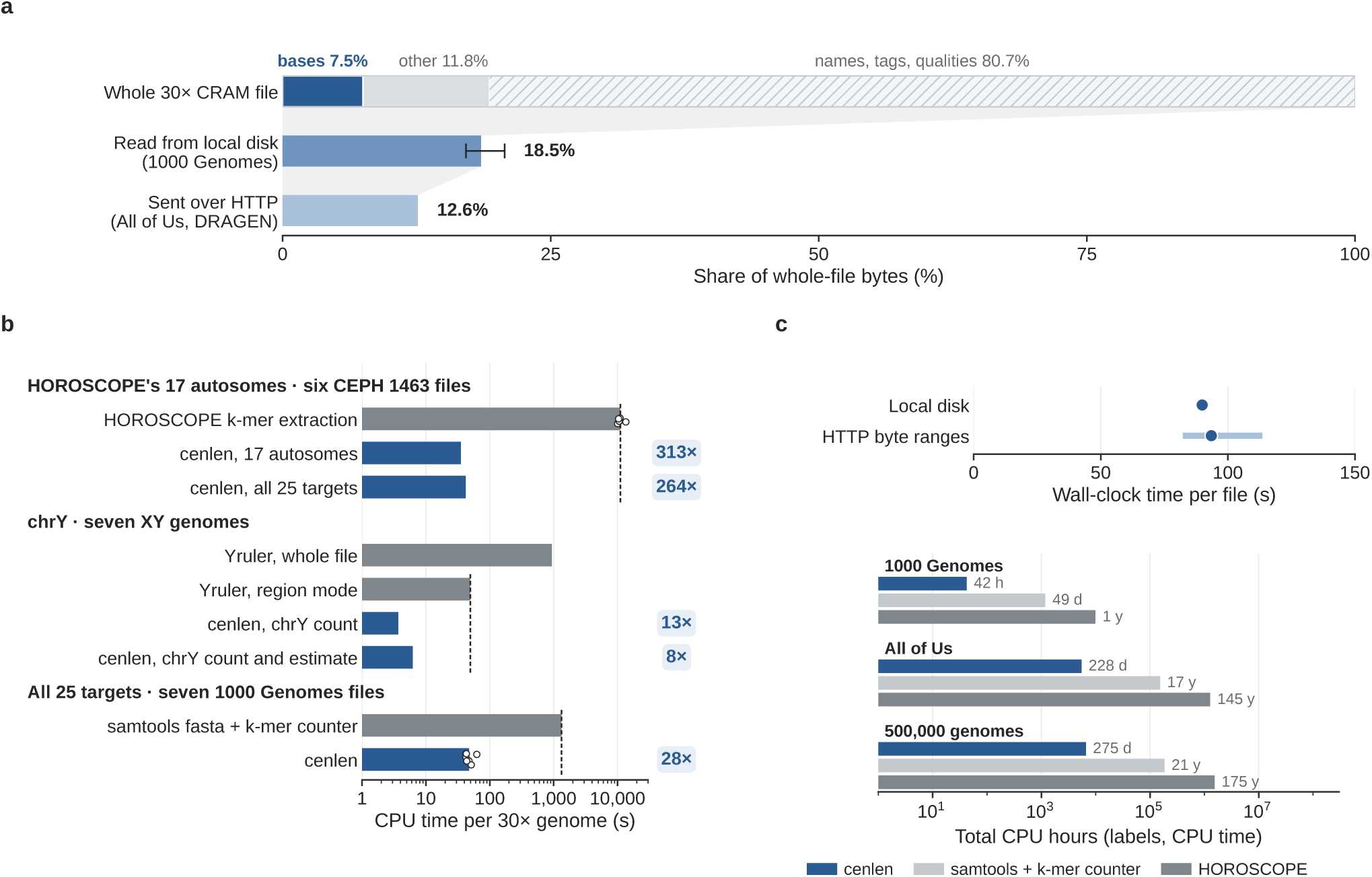
Targeted decoding reduces data retrieval and compute. (a) Share of whole-file bytes: compressed byte composition of a 30× 1000 Genomes CRAM file (sequence bases 7.5%; other 11.8%; read names, alignment tags and base qualities 80.7%, not decoded), bytes read by cenlen from local disk (18.5% in HG01192, 7,206 of 70,998 containers; line, 17.1–20.7% across seven files) and bytes sent by HTTP byte-range requests from *All of Us* DRAGEN CRAM files (12.6% of a whole-file copy). (b) CPU time per 30× genome on matched compute nodes at three task scopes: HOROSCOPE’s 17 autosomes on six CEPH 1463 files, chrY on seven XY genomes, and all 25 targets on seven 1000 Genomes files (reference pipeline, samtools fasta piped into a k-mer counter). Dots, per-file values; dashed lines, the reference row of each scope (grey bars, other tools); right, speed-up of cenlen over that reference. cenlen peak memory was 0.65 GB. (c) Top, wall-clock time per *All of Us* CRAM file by HTTP byte-range requests (median; bar, 10th–90th percentile) and from local disk (derived from the measured ratio). Bottom, CPU hours to profile the 3,202 high-coverage 1000 Genomes genomes, the 414,830 *All of Us* short-read genomes and a projection for 500,000 genomes (UK Biobank scale) at the measured per-genome CPU time of cenlen (47.5 s), the reference pipeline and HOROSCOPE; labels, total CPU time (h, hours; d, days; y, years).

Estimating all chromosomes took 42 CPU seconds per genome. for HOROSCOPE’s 17 autosomal arrays, cenlen took 35 CPU seconds against 11,060 for HOROSCOPE’s whole-file k-mer extraction, a 313-fold reduction. for the Y chromosome, counting and estimation took 6.2 CPU seconds, 8-fold less than Yruler’s single-core region mode (49.6 s).

### Global diversity of human centromere lengths across 2,557 unrelated genomes

Across all 24 chromosomes, the vast majority of variation in centromeric array length resides within, rather than between, human populations (fig. 5; Supplementary Table S14; Supplementary fig. S3). In the 2,557 unrelated founders of the high-coverage 1000 Genomes Project, the haplotype-scale share of variance between superpopulations had a median of 0.045 across the 22 autosomal arrays, below the 0.068 of 114,338 LD-pruned common SNVs, and ranged from 0.005 (chr3) to 0.171 (chr21). Population medians also covaried across chromosomes (fig. 5b; share 0.157 for the genome’s mean log length; Supplementary Table S15).

**Figure 5:**
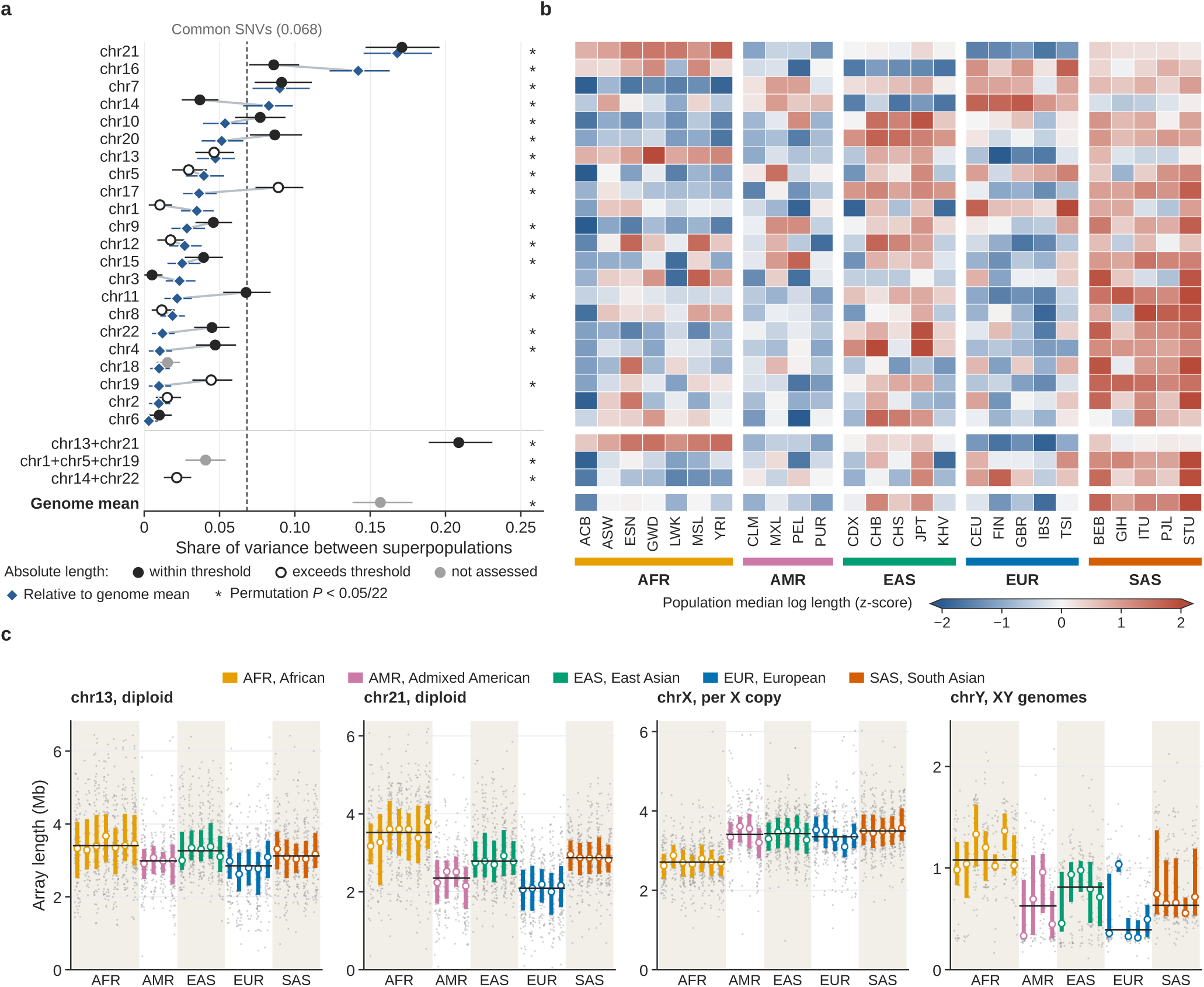
Global diversity of centromere array length in 2,557 unrelated genomes of the 1000 Genomes Project. (a) Share of variance between superpopulations on the haplotype scale, *V*_super_/(*V*_super_ + *V*_pop_ + 2*V*_within_), with 95% bootstrap intervals. Circles, absolute log length: filled, within the cross-ancestry residual-bias threshold (superpopulation differences in mean out-of-fold training residuals below half the range of superpopulation medians); open, exceeding it; grey, not assessed. Diamonds, log length relative to the genome mean of the other 21 autosomal arrays; lines join each array’s two shares. Autosomes, ordered by relative share, precede the three shared-HOR family totals and the genome mean (mean log length of the 22 autosomal arrays). Dashed line, share across 114,338 common SNVs (0.068); asterisks, permutation *P <* 0.05/22 (absolute length; 10,000 permutations of whole populations). (b) Population medians of log length standardized across the 26 populations (z-scores capped at ±2; 10 of 676 cells beyond), rows as in (a); AfR, African; AMR, admixed American; EAS, East Asian; EUR, European; SAS, South Asian. (c) Array length by population for chr13 and chr21 (diploid totals), the X array per copy (XX totals halved) and the Y array of 1,268 XY genomes: grey points, genomes; bars and dots, population interquartile ranges and medians; dark lines, superpopulation medians; axes end at 1.06 times the 99.8th percentile (2–5 genomes per panel lie above). All 24 chromosomes are shown in Supplementary fig. S3.

Because normalization against the mean of the other autosomes eliminates a shared genome-wide scaling factor (raising the median correlation with assembled lengths to 0.992; Methods; Supplementary Section S5), we evaluated both absolute and relative array lengths (fig. 5a). Three arrays diverged between populations more than common SNVs: chr21 was longer in the African superpopulation (median diploid total 3.52 Mb against 2.09–2.88 Mb elsewhere); chr7 was shorter in the African superpopulation (5.64 Mb against 6.71–6.88 Mb); and chr16 was shorter in the East Asian superpopulation (3.29 Mb against 3.52–3.93 Mb).

### X-array length associates with *MED14* splicing in 730 lymphoblastoid cell lines

Whether variation in centromere array length influences downstream gene regulation or RNA processing remains largely unexplored at population scale. To assess the possible role of array length in gene regulation, we performed association analyses between array length and gene expression and splicing phenotypes derived from 730 donors in LCL cells [17]. We observed that 23 genes were associated with array lengths (*q <* .05), and that 11 chromosomes had at least one associated array length-molecular trait pair after stringent sensitivity analyses (Supplementary S10).

Because array lengths may be associated with genotypes of SNPs near the array, we then performed a sensitivity analysis conditioning on the three strongest proximal SNPs. We found that 13 of these associations persisted after this adjustment (fig. 6a,c).

**Figure 6:**
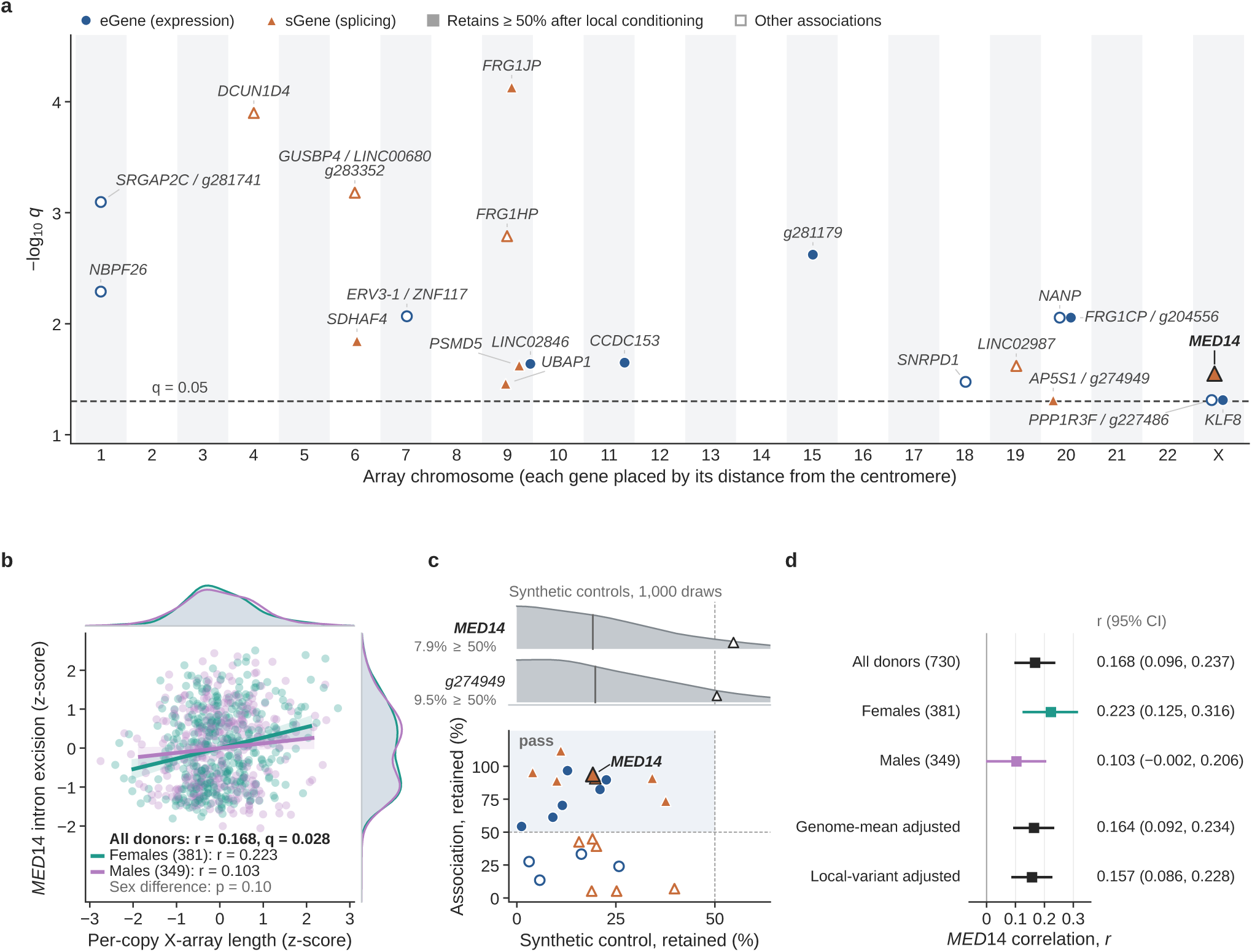
X-array length associates with *MED14* splicing in 730 MAGE lymphoblastoid cell lines. (a) Associations of array length with expression (eGenes, circles) and intron splicing (sGenes, triangles) at *q <* 0.05 for genes on the array’s chromosome, by array and, within each array, by distance from the centromere (28 genes or introns on 11 arrays; introns of one splicing cluster and coincident genes share a point). filled markers denote associations that pass the robustness checks (Supplementary Section S10) and retain at least half of their correlation after conditioning on the three local variants most strongly associated with the phenotype while a synthetic tag-variant control does not; for two associations, including *MED14* (outlined), the control was evaluated over 1,000 draws (c). g followed by six digits abbreviates the Ensembl identifier ENSG00000 followed by those digits. (b) Normalized *MED14* intron excision against per-copy X-array length (inverse-normal within sex) in 381 female and 349 male donors, both adjusted for five genotype principal components and 15 PEER factors, with per-sex regression lines, 95% confidence bands and marginal densities; sex difference by fisher *z*. (c) Bottom, fraction of correlation retained after conditioning on three local variants for the 23 robust associations against that of their synthetic controls; the shaded quadrant marks associations that pass; outlined, the two associations with an inconclusive single control draw, at the median of 1,000 draws. Top, the distribution of their 1,000 control draws (line, median; open triangle, the primary-scan draw; percentages, share of draws retaining at least 50%); *MED14* retained 94% against a control median of 19%. (d) Correlation of X-array length with *MED14* splicing (fisher-*z* 95% confidence intervals) across all donors, by sex, and after adjustment for genome-mean array length or for the three local variants.

We observed that chromosome X array length was associated with splicing of *MED14* (*r* = 0.17; *q* = 0.028 across 388 X-linked genes; fig. 6b; Supplementary Table S16). *MED14* encodes a Mediator subunit whose inactive-X allele undergoes reactivation upon *XIST* loss [18, 19] and is ∼17 Mb from the chr X centromere. The association had the same direction in both sexes (*r* = 0.22 in females and 0.10 in males; fisher *z* test of homogeneity, *p* = 0.10) and was unchanged after adjusting for genome-wide mean array length (*r* = 0.164) or for the three local variants most strongly associated with the intron (*r* = 0.157) (fig. 6c,d).

## Discussion

Alpha-satellite higher-order repeat (HOR) arrays are among the most variable regions of the human genome [1, 20], yet megabase-scale length measurements have been restricted to the few hundred individuals with complete long-read assemblies [3, 4]. In this study, we developed cenlen, an approach that accurately estimates the active array length of every human centromere directly from short-read sequencing data in just under one CPU-minute per genome (Table 1). cenlen thus enables population-scale studies of centromere array length variation and its relationship to molecular, cellular, and organismal phenotypes.

Two methodological innovations distinguish cenlen from previous repeat-profiling and k-mer counting ap- proaches. Algorithmically, prior short-read estimators rely on chromosome-unique sequence tags [2, 13], which are insufficient in acrocentric and repeat-sharing chromosomes where concerted evolution has homogenized repeat units [15, 21]. *cenlen* compares normalized k-mer counts with profiles derived from assembled centromeric arrays. This allows the model to account for differences among arrays of the same chromosome and for repeat sequences shared by different chromosomes. Second, cenlen redesigns the computations by replacing whole-file streaming with index-guided retrieval and efficient k-mer counting. Both design principles could naturally extend to other multicopy or paralogous loci where reads hard to map uniquely, including ribosomal DNA clusters [22], variable number tandem repeats [23], and segmental duplications [24], provided that assembled reference panels represent their structural diversity. We note cenlen’s efficiency optimizations is likely suitable for other k-mer based approaches [13, 25].

As with non-centromeric genetic variation, we find that most variation in centromere active array length lies within populations as opposed to between populations. However, several arrays, most notably on chr21, chr7 and chr16, differed between superpopulations more than the average common SNPs, and the chr21 array was longest in the African superpopulation. Centromeric arrays can expand rapidly through unequal exchange [3, 4] and are embedded in long, recombination-suppressed haplotypes that can retain archaic lineages [26, 27]. These features may contribute to the observed population differences in array length.

To investigate the extent to which array length variation might be functional, we linked cenlen array length estimates to genes expression and splicing profiles. We found several molecular phenotypes associations, including the alternative splicing of *MED14* in chromosome X. Recent studies have linked *MED14* reactivations to the inactive X upon *XIST* loss [18]. One explanation is that megabase-scale repeats act as heterochromatic sinks that titrate chromatin factors [28, 29].

Several considerations outline directions for future work. first, short-read estimates for autosomes and for chrX in XX genomes are still unphased diploid totals; dissecting homologue-specific and parental effects will require pedigree transmission analysis or statistical phasing against long-read panels. Second, resolving chr13 and chr21 individually remains challenging because their active arrays share 96–97% of their k-mer copies, which widens their prediction intervals. Expanding pangenomic reference panels with broader haplotype diversity should provide the fine-scale profile distinctions needed to separate these two acrocentric chromosomes cleanly. Third, deconvolution can assign weight largely to array profiles represented in the reference panel, and rare or population-restricted HOR variants absent from current assemblies [4, 30] may bias estimates; incorporating upcoming pangenome releases with broader global diversity and representation [31] will reduce the bias.

In conclusion, cenlen uses pangenome-derived reference panels and algorithmic innovations to estimate cen- tromeric active array lengths. We anticipate that our work will enable the standard inclusion of centromere array lengths in large-scale human genetic association studies.

## Methods

### Overview

cenlen estimates the active alpha-satellite array length of every human chromosome from a single short-read alignment file (fig. 1b,c; Table 1). Estimates for autosomes and for the X chromosome of XX genomes are diploid totals, *L*_diploid_ = *L*_1_ + *L*_2_, whereas the X estimate of XY genomes and the Y estimate are single-copy lengths. The algorithm counts a fixed query of canonical 31-mers across targeted genomic intervals, normalizes counts against single-copy 31-mers tallied within the same intervals, converts normalized abundances into lengths using models calibrated on complete assemblies, and computes an empirical prediction interval for each estimate. Chromosomal sex, inferred directly from these k-mer abundances, determines which X-chromosome model is evaluated and enables the Y- chromosome estimate.

### Assembled arrays and training genomes

We used assembled active arrays from HGSVC3 [8], the Human Pangenome Reference Consortium [32] and Platinum Pedigree [16], as annotated by Gao et al. [4]. for each haplotype, the annotated array length served as the reference length (Supplementary Section S3). Of 310 genomes with assembled arrays, 282 also have 30× Illumina reads from the New York Genome Center [33]. We partitioned genomes by pedigree family into a validation set of 62 sequenced genomes (61 of which have at least one completely assembled target array) and a training set of 248 genomes (220 sequenced and 28 assembly-only; Table 2). We estimated all model parameters (k-mer densities, reference panels, calibration factors and prediction-interval residuals) within the training partition. for each genome, the reference length of a target array was the sum of its active-array haplotype lengths. We included a genome in model fitting for a target only when all required haplotypes were completely assembled. We also evaluated cenlen in six members of the CEPH 1463 pedigree sequenced at the same center. Their genomes were not included in the training or initial validation sets (see Evaluation).

**Table 2:** Genomes used in this study. Pairs, genome–array pairs with a completely assembled reference array.

| Cohort | Genomes | Use |
|---|---|---|
| Assembled genomes | 310: 282 with 30× CRAM files, 28 assembly-only | Reference lengths and panels |
| Training partition | 248: 220 sequenced, 28 assembly-only; k-mer selection in 105 | k-mer sets, models, calibration, intervals |
| Validation partition | 62 sequenced; 61 with a complete array (566 pairs) | Held-out accuracy and coverage |
| Nested cross-validation | 282 sequenced; 280 with a complete array (2,745 pairs) | Accuracy with every fitting step refitted |
| CEPH 1463 | 6 genomes with Platinum Pedigree assemblies (83 lengths) | Accuracy and comparison with HOROSCOPE |
| 1000 Genomes, high coverage | 3,202 genomes; 427 trios; 308 father–son pairs; 2,557 unrelated founders<sup>a</sup> | Transmission, sex classification, population diversity |
| MAGE | 730 lymphoblastoid cell lines (381 female, 349 male) | Expression and splicing associations |

<sup>a</sup>After excluding six genomes failing depth quality control and three discordant with pedigree sex, and pruning one genome from each of 29 related pairs among 2,583 founders (Supplementary Section S9).

### The 31-mer query

for each chromosome, candidate k-mers were the canonical 31-mers of its annotated active arrays. for chromosomes with a chromosome-specific array, we retained a candidate 31-mer if it was rare in other centromeres and in regions of the same centromere outside the active array. We also required the combined copy number of its one-mismatch (Hamming distance 1) neighbours to remain below a threshold. The 31-mer had to occur in at least a predefined fraction of selection haplotypes, both overall and within each continental ancestry group. This procedure yielded a chromosome-specific set *S_c_* of 1,025–3,949 k-mers (Supplementary Section S1; Supplementary Tables S3 and S4). Some chromosomes share homologous HORs [15]. We grouped chr13 with chr21 and chr14 with chr22. We also fitted chr1, chr5 and chr19 jointly with chr16. Within-family pairs share 41–97% of k-mer copies, compared with at most 34% across other pairs (fig. 2b). for each family, we retained 31-mers that occurred in at least 5% of training arrays for at least one family member and were rare outside the family. These k-mers formed the shared family set *S_g_* . The indexed binary query table contains 1,021,335 canonical 31-mers, of which 915,865 enter the active estimation models (comprising chromosome-specific sets, family sets, the autosomal depth panel and the two chrY sets) and are queried in a single pass (Supplementary Section S2).

### CRAM container filtering and byte-stream parsing

The extraction module is written in C and linked against htslib [34]. It parses an indexed CRAM file in a single pass over a fixed set of 3,432 targeted genomic intervals: the GRCh38 centromeric bands with flanks, annotated heterochromatin, the acrocentric short arms, the whole chrY contig, the unplaced, decoy, alternative, HLA and EBV contigs, and the unmapped reads (Supplementary Section S2; Supplementary Table S10). Within the containers that overlap these regions it decodes only read flags, reference identifiers and bases [35], and retains all primary records, including mapped, unmapped and duplicate-flagged alignments, thereby counting centromeric reads irrespective of their alignment coordinates. Duplicate-flagged reads are retained for calibrations (Supplementary Section S12).

K-mer counting is coupled to slice decoding, which removes intermediate serialization and inter-process commu- nication. Each read sequence is pre-screened in 16-base blocks against a blocked two-bit Bloom filter of all 16-base substrings of the query k-mers; exact lookups are restricted to the 31-mer windows that contain a filter hit. for reads identical to the reference sequence, k-mer occurrences are aggregated once per slice from the reference sequence. Container slices are processed by a dedicated slice decoder, with htslib for the remaining slice types. The program counts every 31-base window containing only A, C, G or T in canonical form [36], producing a count *n_k_* for each query k-mer.

### Canonical k-mer counting and depth normalization

To establish baseline sequencing depth, we curated a reference panel *K*_pos_ comprising 129,274 single-copy autosomal 31-mers situated within the targeted genomic intervals (Supplementary Section S2). The per-copy coverage *d* is defined as

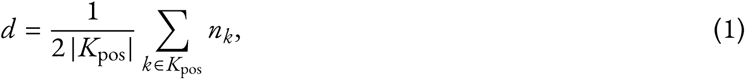

from which the normalized k-mer abundance is computed as *y_k_* = *n_k_*/*d*. Because we calculate *d* from the same containers as the array counts, changes in sequencing depth and in the fraction of reads retrieved affect *n_k_* and *d* similarly. Dividing *n_k_* by *d* therefore largely removes these effects from *y_k_*.

### Statistical framework for length estimation

four models convert normalized counts into lengths: the specific-k-mer count, the family-mean mixture, the reference panel fitted by non-negative least squares, and the Y-specific count described below.

We use the same framework to describe k-mer counts in the three models for autosomes and chromosome X. An assembled array *a* with *N_ak_* occurrences of k-mer *k* over its length ℓ*_a_* has the per-base-pair profile *P_ak_* = *N_ak_*/ℓ*_a_*. If a genome’s arrays are represented by lengths *β_a_* ≥ 0 (base pairs) assigned to profiles *P_a_*_·_, the normalized count of *k* has expectation E[*y_k_*] = Σ*_a_ β_a_ P_ak_*, and Poisson read sampling at per-copy depth *d*, *n_k_* ∼ Poisson (*d* E[*y_k_*]) , gives Var(*y_k_*) ≈ E[*y_k_*]/*d*. The lengths are estimated by non-negative weighted least squares with weights approximating the inverse variance, and each member’s estimate is the calibrated sum of its coefficients:

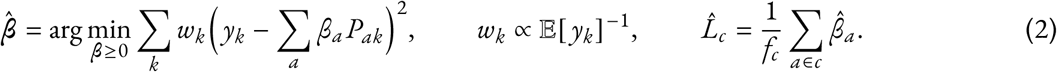

The models differ in the columns *a*: one mean-density column for the specific-k-mer count, for which Eq. 2 reduces to the ratio estimator below; one mean-density column per member for the family-mean mixture; and one column per assembled training array for the reference panel. E[*y_k_*] in the weights is replaced by its value at typical array lengths, so the weights do not depend on the genome being estimated.

## Chromosome-specific sets

For a chromosome *c* quantified by its set *S_c_* (chr18 and chrX), the expected number of set occurrences per base pair of assembled array is

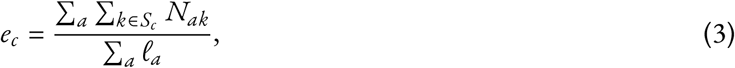

where *N_ak_* is the number of occurrences of k-mer *k* in training array *a* of chromosome *c*, ℓ*_a_* is that array’s length, and both sums run over the training arrays of *c*. The length estimate is

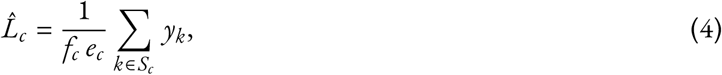

with *f_c_* the calibration factor (below).

### Family mixture

for chr13 and chr21, we model the counts of the shared family k-mer set as a mixture of contributions from the two chromosomes. for chromosome *c*, the model uses the mean density *D_ck_* = *_a_ N_ak_*/Σ *_a_* ℓ*_a_* across its training arrays, scaled by its unknown length *L_c_*.

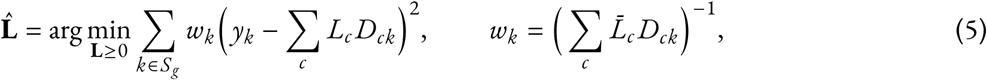

where 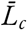 is twice the mean haplotype length of member *c*, so that *w_k_* is proportional to the inverse of the Poisson variance expected at the members’ mean diploid lengths, 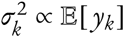. With two members the problem is solved exactly by enumerating the active sets of the non-negativity constraints. The estimated array length of member *c* is 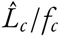

#### Panel model

Arrays of one chromosome differ in HOR composition, so a uniform mean density systematically underestimates or overestimates atypical array variants. The reference panel replaces the mean density by the k-mer profiles of individual assembled arrays. A group *g* consists of either one chromosome and its set *S_c_* or one family of chromosomes. The 13 single-chromosome groups and two family groups comprise 19 panel targets. The family groups contain chr14 and chr22, and chr1, chr5 and chr19 fitted jointly with chr16 (Supplementary Table S4). A family group uses the 10,000 k-mers of its family set with the largest member contrast, 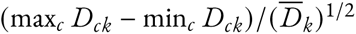, where 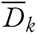 is the mean of *D_ck_* over members.

The panel of group *g* consists of the training arrays of its members, 156–1,174 arrays per group (Supplementary Table S4). for each array *a* from chromosome *m*(*a*), the panel includes its per-base-pair profile *P_ak_* = *N_ak_*/ℓ*_a_* over the group’s k-mers. An array whose density Σ_k_*N_ak_*/ℓ is below half the median density of its member’s arrays is excluded, which removes arrays with truncated or misannotated sequence. Let 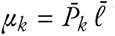 be the panel’s mean profile times its mean array length, the count of k-mer *k* expected from a typical array; k-mers with *μ_k_* = 0 are filtered out, and the rest are weighted by *w_k_* = max(*μ_k_*, 1)^−1^, which approximates the inverse variance of normalized counts under Poisson read sampling at fixed per-copy depth,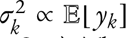 (validated against nine alternative weighting schemes across all 21 autosomes in Supplementary Section S11). The coefficients

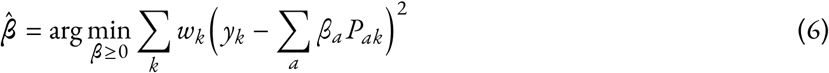

are obtained by non-negative least squares on rows scaled by 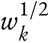 (Lawson–Hanson algorithm as implemented in SciPy [37]). Each coefficient represents the length, in base pairs, assigned to the composition of one panel array. for chromosome *c*, we sum the coefficients for its arrays and divide by *f_c_* to obtain the length estimate,

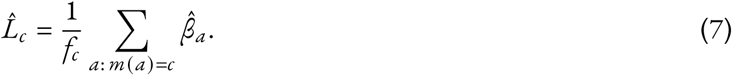

#### Model assignment

The reference panel is the default model for autosomal arrays because the assembled haplotype profiles represent variation in HOR composition and sequences shared across chromosomes. A simpler model is retained where the assembled training set cannot support a multi-array basis (chr18, *n* = 13 arrays), where near-complete cross-chromosome homology and assembly-only outlier haplotypes destabilize panel weights (the chr13– chr21 family; Supplementary Section S4), and for chrX, which is modeled by its chromosome-specific k-mer set. An automated rule applied to training cross-validation predictions implements these criteria and is repeated within every outer split of the nested cross-validation (Supplementary Section S5).

#### Calibration factors

for each target, uncalibrated estimates from short reads differ from assembled lengths by a near-constant factor, reflecting sequencing errors, mappability and finite read sampling. We calculated the calibration factor *f_c_* as the median ratio of the uncalibrated estimate to assembled length across training genomes with complete arrays. Calibration leaves correlations unchanged and lowers the pooled validation median relative error from 3.2% to 2.7% across the 24 calibrated targets. for reference panels, each training genome is evaluated after excluding haplotypes from its own pedigree family, mirroring out-of-sample evaluation. The X chromosome has two calibration factors, one from XX genomes (diploid total: both arrays) and one from XY genomes (one array). The calibration factors lie between 0.936 and 1.090 (median 0.992; Supplementary Table S4).

#### Sex classification

The sex classification uses 3,290 single-copy k-mers of the male-specific region of chrY. Their mean count *u_Y_* , relative to the autosomal depth, gives *R_Y_* = *u_Y_* /*d*; the genome is classified as XY (M) when *R_Y_* ≥ 0.365, XX (f) when *R_Y_ <* 0.05 and unclassified (U) otherwise. This classification determines which X target applies and whether chrY is estimated; for a genome of unclassified sex, cenlen reports both X estimates and the Y estimate.

#### The Y array

The active array of chrY, DYZ3, is a single-copy array susceptible to mosaic loss of chromosome Y (mLOY) in a fraction of cells, so its k-mer count is normalized by the single-copy reference k-mers in the male-specific region of the Y chromosome (MSY), as used for sex classification:

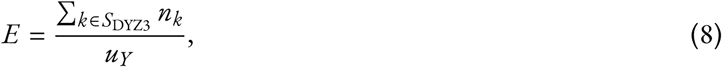

where *u_Y_* denotes the mean abundance of the MSY normalization k-mers, so that *E* is measured per copy of the Y chromosome, making the estimate invariant to mLOY. The 3,816 k-mers of *S*_DYZ3_ occur at a constant density along assembled DYZ3 arrays, so *E* is proportional to length, and calibration is a zero-intercept linear model, 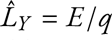with *q* (normalized counts per Mb) the median of *E*/*L* over the 59 training XY genomes with an assembled array (Supplementary Section S7).

#### Prediction intervals

Each estimate includes 90% and 95% prediction intervals derived from empirical quantiles of out-of-fold residuals, a cross-validation analogue of split-conformal prediction whose coverage is evaluated empirically. for target *o*, the residuals are the log prediction errors 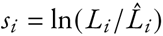, the log ratio of assembled length to the cross-validated prediction for training genome *i* (five-fold cross-validation with folds formed by family; for chrY, the training XY genomes, with *q* refitted within each fold). With *n* residuals and miscoverage *α* ∈ {0.10, 0.05}, let *l* = ⌊(*n* + 1) *α*/2⌋ and *u* = ⌈(*n* + 1)(1 − *α*/2)⌉. When 1 ≤ *l < u* ≤ *n*, the interval is

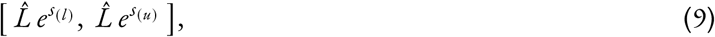

with *s*_(1)_ ≤ · · · ≤ *s*_(_*_n_*_)_ the order statistics of the target’s own residuals. for targets whose training sample size is insufficient for direct order-statistic quantiles (*n <* ⌈(2 − *α*)/*α*⌉), standardized residuals are pooled within the corresponding model class (reference panels; the chr13–chr21 family; and specific-k-mer counts): each residual is standardized as 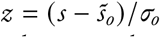 where 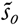is the leave-one-out median residual and *σ_o_* is 1.4826 times the leave-one-out median absolute deviation shrunk toward the class median scale with weight *n_o_*/(*n_o_* + 10). The prediction interval is then 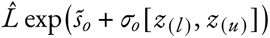 using quantiles *z*_(_*_l_*_)_ , *z*_(_*_u_*_)_ of the pooled distribution. Cross-target standardized pooling serves as a small-sample empirical approximation for chr18 (90% and 95% intervals) and chr17 (95% interval), whereas all other targets use target-specific empirical quantiles (*n* = 22–169 per target). On synthetic residuals of the measured sizes, mean coverage was 0.89–0.91 at 90% and 0.94–0.96 at 95%.

#### Relative array length

Errors in array length estimates have a component shared across chromosomes within each genome. Across training genomes, out-of-fold residuals for different targets had a median pairwise correlation of *r* = 0.77, whereas assembled lengths of different autosomes had a median pairwise correlation of *r* = 0.001 (Supplementary Section S5; Supple- mentary Table S9). This component acts as a genome-level scaling factor and is eliminated in the relative length of autosomal array *c*, 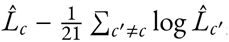, the log estimate minus the genome’s mean log estimate over the other 21 autosomal arrays. In the training genomes, relative lengths lowered the standard deviation of the log error from 4.3% to 2.6% and raised the median correlation with assembled values from 0.972 to 0.992, with higher correlation on all 22 autosomes (Supplementary fig. S2b,c).

#### Implementation

The core extraction module is a statically linked C program. The estimator is a Python program using NumPy, SciPy and pandas. It takes the count vector and fixed model files as input, including k-mer positions, set densities, panel profiles as integer occurrence counts with array lengths, calibration factors and interval factors. Targeted k-mer counting and downstream length estimation execute as two modular stages. In addition to local CRAM parsing, an HTTP-enabled build retrieves index metadata and target containers directly from cloud object storage via byte-range requests.

#### Evaluation

#### Training, validation and nested cross-validation

five-fold cross-validation across pedigree units of the 248 train- ing genomes provided training accuracy metrics and prediction-interval residuals, and the released model parameters were evaluated on the 62 validation genomes (61 with at least one completely assembled target array). To obtain an unbiased generalization estimate across the full model-selection and calibration workflow [38], we additionally performed five-fold family-stratified nested cross-validation across all 282 sequenced genomes (grouping connected pedigree and kinship components with KING *>* 0.05), repeating candidate filtering, automated model assignment, calibration and interval construction within each outer training split before predicting held-out outer-fold genomes (Supplementary Section S5). We also evaluated six CEPH 1463 genomes sequenced at the same center. for each target, we report sample size, Pearson *r*, median absolute relative error (MedRE), RMSE and prediction-interval coverage; pooled coverage is reported with a cluster-bootstrap confidence interval over pedigree units to account for within-genome and familial error correlation.

#### Families

Transmission in parent–offspring trios of the high-coverage 1000 Genomes cohort provides an assembly- independent check. for each target, the transmission slope *R* is the regression coefficient of the child’s estimate on the mid-parent estimate, corrected for the correlation between spouses, with a bootstrap of 2,000 resamples of families. *R* = 1 under error-free measurement of a strictly Mendelian trait; non-inherited measurement noise attenuates *R* toward zero, whereas haplotype-correlated estimation bias co-inherited with the array can elevate *R* above unity (Supplementary Section S8).

#### Comparison with existing methods

We used six CEPH 1463 pedigree members with 1000 Genomes CRAM files (NA12877, NA12878, NA12889, NA12890, NA12891 and NA12892; two parent–offspring trios) to compare estimates with Platinum Pedigree reference array lengths [16]. None of these genomes were included in the cenlen training or validation sets or among HOROSCOPE’s 272 training samples [13]. We compared cenlen and HORO- SCOPE across all genome–array pairs with estimates from both methods (Supplementary Section S6). We doubled HOROSCOPE’s per-copy HOR length to match cenlen’s diploid sum and reported array-averaged median absolute error, mean absolute error, RMSE and bias. We also compared cenlen with a linearly recalibrated HOROSCOPE baseline fitted separately for each array using 59 of HOROSCOPE’s training genomes. To assess the influence of related individuals, we repeated the comparison leaving out each individual and each trio among the six pedigree members. for chrY, we compared cenlen with Yruler [12] across the validation XY genomes and across all 70 XY genomes with complete assemblies. Supplementary Section S6 details the comparison and benchmarking protocols.

#### Population-scale profiling

cenlen estimated every target in the 3,202 high-coverage genomes of the 1000 Genomes Project [33]. Six genomes failed the sequencing-depth filter and three the sex-classification filter; the primary sample comprised the 2,557 pedigree founders that remained after removal of one genome from each pair with KING kinship above 0.0884 [39]. for each array, a nested analysis of molecular variance [40] partitioned the variance of log length into components between the five superpopulations (*V*_super_), between populations within superpopulations (*V*_pop_) and within populations (*V*_within_), with 95% intervals from 1,000 bootstrap resamples within populations; *V*_super_ was tested by permuting whole populations among superpopulations (10,000 permutations) and *V*_pop_ by permuting individuals among the populations of each superpopulation. Because each estimate is a diploid total (*L*_1_ + *L*_2_), the share between superpopulations on the haplotype scale is*V*_super_/(*V*_super_ +*V*_pop_ + 2*V*_within_); the same estimator applied to 114,338 common single-nucleotide variants (SNVs), pruned for linkage disequilibrium, gives the reference for genome-wide variation, both as the ratio of summed components (the average common SNV) and as the distribution of single-SNV shares. To assess potential ancestry-associated calibration shifts, mean out-of-fold training residuals were compared across superpopulations against a threshold equal to half the inter-superpopulation median range. Each array was analysed in absolute and in relative length.

#### Technical replicates

The 103 short-read alignments of the SEQC2 reference pair HCC1395 and HCC1395BL [41, 42] (BioProject PRJNA489865) were read over fTP for the target regions, which held a median 9.2% of each file’s records. Their reference build lacks the alternative and HLA contigs; in a whole-file comparison of one library, estimates changed by a median 0.13% (Supplementary Section S12).

#### Benchmarks

Programs were timed on shared nodes with Intel Xeon Gold 6430 processors, each comparison on the same files and node. cenlen and Yruler ran single-threaded and pinned to one core, HOROSCOPE’s k-mer extraction ran in its workflow on eight threads, and a reference pipeline, samtools fasta [43] piped into a k-mer counter, ran with two and four threads. CPU time is user plus system time summed over threads, and memory is the peak resident set size. Ablation benchmarks, in which single features were disabled and each configuration was run twice in mirrored order, quantified the speed contributions of container selection, data-series selection, decode-coupled counting, Bloom prefiltering and per-slice reference counting under identical query tables (Supplementary Section S6; Supplementary Table S13). for equal task scope, cenlen’s target list was also restricted to HOROSCOPE’s 17 autosomes or to the chrY contig (Supplementary Section S6). Retrieval from cloud storage was measured in the *All of Us* Researcher Workbench [9] on DRAGEN-aligned CRAM files (Supplementary Section S8).

## Code availability

cenlen is written in C and Python and distributed under the MIT licence at https://github.com/YikaiDong-g it/cenlen; the source code and test data used in this study will be archived at Zenodo upon publication.

## Acknowledgements

This study used data from the *All of Us* Research Program’s Controlled Tier Dataset (Curated Data Repository version 8), available to authorized users on the Researcher Workbench. The *All of Us* Research Program is supported by the National Institutes of Health, Office of the Director. We gratefully acknowledge *All of Us* participants for their contributions, without whom this research would not have been possible. We thank the Human Pangenome Reference Consortium, the Human Genome Structural Variation Consortium, the 1000 Genomes Project and the New York Genome Center for making their data publicly available.

## Data availability

Reads of the high-coverage 1000 Genomes Project [33] are available from the European Nucleotide Archive (PR- JEB31736 and PRJEB36890), with CRAM alignments hosted at s3://1000genomes/1000G_2504_high_co verage/. Centromeric array sequences and CenMAP annotations for the HGSVC3, HPRC and Platinum Pedi- gree assemblies are available on figshare [4] (doi:10.6084/m9.figshare.29921990). Platinum Pedigree short reads [16] are hosted at s3://platinum-pedigree-data/. MAGE transcriptomic data [17] are archived at Zenodo (doi:10.5281/zenodo.10535719). SEQC2 alignments of HCC1395 and HCC1395BL [41, 42] are available from the NCBI fTP site (ftp://ftp-trace.ncbi.nlm.nih.gov/ReferenceSamples/seqc/Somatic_Mutation_WG/), with reads under BioProject PRJNA489865. *All of Us* data are accessible to authorized researchers via the Researcher Workbench. Supplementary Data 1 lists the analytical partition of each genome, and Supplementary Data 2 reports all estimates for the validation and CEPH 1463 genomes alongside assembled reference lengths and prediction intervals.

## Contributions

Y.D. and J.W. conceived the study. Y.D. developed the methodology, implemented the software and performed the formal analyses, validation and visualization. J.W. provided resources and supervised the study. Y.D. wrote the original draft. Y.D. and J.W. reviewed and edited the manuscript. All authors read and approved the final manuscript.

## Supplementary material

### S1 Genomes and k-mer sets

Supplementary Data 1 lists the analytical assignment of each genome. We assigned related individuals to the same partition. Among 310 reference-assembled genomes, 282 had New York Genome Center 30× CRAM alignments and 28 were assembly-only training genomes. We partitioned pedigree-connected family units outside the k-mer selection subset into training and validation sets. The validation set comprised 62 sequenced genomes, 61 of which had at least one completely assembled target array. We prioritized family units containing complete chr17 and chr18 pairs. The training set comprised 248 genomes. Chromosome-specific k-mer sets were selected across haplotypes of 105 training genomes according to Table S3; shared-family sets retain 31-mers with prevalence ≥ 0.05 among training arrays of at least one family member and, on every non-member chromosome, mean copy number ≤ 0.005 of the family mean and prevalence ≤ 0.05 (Table S4).

### S2 Target regions, retrieval and normalization

The target regions (Table S10) are released as a BED file of 3,432 intervals on the reference of the 1000 Genomes CRAM files (GRCh38 with decoy and HLA sequences); for a CRAM file aligned to another reference, cenlen resolves the list against the file header. Targeted extraction retrieves every overlapping CRAM container from its byte offset to the subsequent container’s offset (7,206 of 70,998 containers in HG01192; 17.1–20.7% of file bytes across seven files); in coordinate-sorted alignments these coalesce into one contiguous byte range per primary chromosome plus one spanning unlocalized, decoy and unmapped records, requiring only a few dozen HTTP range requests per file. Normalization k-mers occur as single-copy sequences in at least a fixed fraction of assembled haplotypes, in T2T-CHM13 and in GRCh38, are absent from non-homologous chromosomes, and lie within 6 Mb of the active array; the 129,274 of 135,278 single-copy k-mers situated within the target intervals constitute *K*_pos_. The binary query table indexes 1,021,335 canonical 31-mers with a bitmask encoding target-set membership, of which 915,865 enter the active estimation models. Technical validation of k-mer counting, array annotations and sample matching is summarized in Table S11.

### S3 Assembled lengths

Array lengths come from the CenMAP annotation of Gao et al. [4]: a monomer is designated active when its name includes the L label, active blocks less than 8 kbp apart are merged, and an array comprises at least five HOR monomers with at least 90% of its length active. HGSVC3 centromeres free of assembly errors and all HPRC and Platinum Pedigree centromeres annotated in that dataset were used. Each haplotype array length equals the summed span of its annotated active HOR blocks; a genome’s reference length is the diploid sum across both haplotypes for autosomes and XX chrX (or the single haplotype span for XY chrX), excluding genomes without complete annotations on all required haplotypes.

### S4 Separation of family members

Within the chr13–chr21 family, active arrays share 96–97% of their k-mer copies and exhibit a wider distribution of short and low-density haplotypes among the 28 assembly-only training genomes (median chr21 length 1.05 Mb versus 1.34 Mb in the 220 sequenced training genomes, including zero-density outlier haplotypes across contrast 31-mers). Incorporating assembly-only training haplotypes into a joint chr13–chr21 panel increased five-fold cross- validated chr21 median relative error from 3.5% to 6.3% and unexplained variance (1 − *r*^2^) by 22%, exceeding the 20% panel-expansion stability bound (chr13 changed by *<* 2% on both metrics); chr13 and chr21 are therefore estimated with the two-member family-mean mixture. Synthetic diploid separation across all three repeat-sharing families is reported in Table S5. Out-of-fold estimation errors of chr13 and chr21 are negatively correlated (*r* = −0.73 in 93 nested cross-validation genomes), whereas their assembled lengths are uncorrelated (*r* = −0.04): the family-mean mixture can assign shared k-mer signal to either chromosome. Errors that shift an equal amount of estimated length from one chromosome to the other cancel in the family total.

### S5 Error structure, model assignment, nested cross-validation and coverage

#### Shared genome-level residual factor

Estimation residuals have a shared genome-level component. Across training individuals, out-of-fold residuals for different targets had a median pairwise correlation of *r* = 0.77 (relative residuals, all 25 targets; 0.81 on log residuals across the autosomes). When array lengths were expressed relative to the genome mean of the other arrays, the median correlation between residuals fell to *r* = 0.20 (fig. S2b); across the autosomes the residual correlation falls to 0.22, the standard deviation of the log error falls from 4.3% to 2.6% and the median correlation with assembled lengths rises from 0.972 to 0.992, with higher correlation on all 22 autosomes and a lower error standard deviation on 20 (fig. S2c). On chromosomes with divergent HOR haplotypes or shared inter- chromosomal repeat cores, replacing specific-k-mer counts and mean-density models with reference panels reduced five-fold training cross-validation RMSE from 0.34, 0.45 and 0.53 Mb to 0.27, 0.14 and 0.27 Mb on chr7, chr10 and chr22, respectively. In nested cross-validation, the median outer-fold RMSE was 0.39, 0.46 and 0.51 Mb under the simpler models against 0.27, 0.15 and 0.26 Mb under the panel (Main fig. 2c).

#### Array length correlations across chromosomes

We assessed correlations between array lengths on different chromosomes and the contributions of estimation error and population structure (Table S9). In the 279 nested cross-validation genomes, assembled log lengths of different autosomes correlate with a median *r* of 0.001, whereas the out-of-fold log errors of the same genomes correlate with a median *r* of 0.78. In the 2,557 unrelated 1000 Genomes genomes, the median correlation of 0.057 between estimated autosomal arrays falls to 0.032 within populations and to 0.008 after subtracting the excess covariance that estimation adds in the cross-validation genomes. Across the 427 trios, whose parents and children were sequenced as separate libraries, twice the covariance between a parent’s array and the child’s array on another chromosome has a median of 0.003 (95% bootstrap interval −0.009 to 0.017), against 0.946 on the same chromosome. The error standard deviation (median 4.4% per array) is small relative to the spread of assembled lengths (standard deviation of log length 12–39% across arrays), so estimation contributes a median of 1.6% of the within-population variance of a single array; the shared error does not average out across arrays and contributes 22% of the within-population variance of a genome’s mean log length, a component that relative lengths remove.

#### Automated model-assignment rule

An autosomal candidate target is assigned the reference panel when three training-set cross-validation criteria hold jointly: (i) cross-validated correlation *r* ≥ 0.80 and |*r* | *<* 0.30 under per- muted reference lengths; (ii) adding training haplotypes from genomes without short-read libraries raises unexplained variance (1−*r*^2^) and median relative error by at most 20%; (iii) across 2,000 pedigree-unit bootstrap resamples, the 95th percentile of the RMSE ratio to the specific-k-mer count or family-mean mixture is at most 1.10 and the 5th percentile of the correlation difference is at least −0.01. All 19 adopted panel targets also keep the Mendelian transmission slope in 1000 Genomes trios above the lower bound of a pedigree simulation (Supplementary Section S8).

#### Nested cross-validation and coverage sensitivity

To evaluate out-of-sample generalization jointly across feature filtering, automated model assignment, calibration and interval construction [38], we performed five-fold family- stratified nested cross-validation across all 282 sequenced genomes (280 with at least one completely assembled target array), grouping pedigree, KING (*>* 0.05) and genomic identity-by-descent relatives into connected components so no related pair crossed outer or inner fold boundaries. Within each outer training fold, family k-mer sets were filtered on outer-training array prevalence (≥ 0.05), top-10,000 contrast k-mers were selected within each inner fold, automated model assignment and empirical interval estimation were executed exclusively on inner cross-validation predictions. We selected the sex-chromosome models for outer-test genomes using the count-derived *R_Y_* classifications. The released assignment retains chr18 (*n* = 13 arrays) at the specific-k-mer count by an explicit small-sample rule; applied without this rule, the automated criteria select the reference panel for chr18 in 4 of 5 outer folds. Across outer folds the reference panel is selected in 5 of 5 folds for 15 autosomes, in 3 or 4 of 5 for chr3, chr4, chr6 and chr11, and in 1 of 5 for chr13 and chr21 (Table S2; fig. S5b). Across all 2,745 held-out genome–array pairs, outer-fold predictions achieved Pearson *r* = 0.907–0.989, median target-specific MedRE of 2.1% (23 of 25 targets ≤ 5%), pooled 90% interval coverage of 91.1% (95% family-cluster bootstrap interval 89.0–92.9%) and pooled 95% coverage of 96.2% (95.0–97.3%; fig. S2a). Restricting evaluation to the 172 genomes (1,587 pairs) whose pedigree and kinship components contain none of the genomes used to select k-mers for single-chromosome targets yielded *r* = 0.901–0.991, median target-specific MedRE of 2.1% and 96.5% pooled 95% coverage (Table S2).

Nested down-sampling of validation read libraries to two-thirds and one-half coverage produced minimal shifts in estimated lengths and interval coverage (Table S8). Supplementary Data 2 reports all estimates for the validation and CEPH 1463 genomes alongside assembled reference lengths and prediction intervals.

### S6 Comparison with HOROSCOPE and benchmark protocol

HOROSCOPE [13] was executed using its published Jellyfish workflow across the six CEPH 1463 CRAM files, doubling its reported per-copy HOR_LENGTH across 17 autosomes (chr1–chr12 and chr16–chr20) to correspond to cenlen’s diploid estimate. The linearly recalibrated baseline fits a separate ordinary least-squares regression of assembled length on twice HOR_LENGTH per chromosome across 59 of HOROSCOPE’s training genomes. The six CEPH 1463 individuals (two parent–offspring trios) formed an independent test set for both methods (Table S6; fig. S5a).

Runtime benchmarks (Table S12) used seven high-coverage 1000 Genomes CRAM files (HG00097, HG00408, HG03732, HG01167, NA18505, HG01192 and HG01243), each counted and estimated twice in counterbalanced order. Scope-restricted runs used identical query tables and parameters while limiting target intervals either to HOROSCOPE’s 17 autosomes (omitting 28 intervals) or to the chrY contig; restricted runs retained 99.988–99.993% of target model k-mer counts, and chrY-restricted counting reproduced full-run chrY estimates within 0.07%. Yruler [12] was benchmarked alongside cenlen’s chrY-targeted run on the same compute node across seven XY genomes (HG03732, HG01167, HG01192, HG01243, NA12877, NA12889 and NA12891), evaluating its targeted region mode (with its published region BED) on one core (two runs per genome in mirrored order alongside cenlen) and on eight threads, as well as single-core whole-CRAM execution (one run per genome). Yruler produced identical estimates on one and eight threads; cenlen’s estimation step on chrY counts required 2.5 CPU seconds per genome. To determine which steps contributed to cenlen’s speed, we benchmarked configurations with individual features disabled on three high-coverage CRAM files (HG00097, HG01192 and NA18505) in two experimental series. We ran each configuration twice in mirrored order using identical query tables (Table S13). Targeted container retrieval and data-series selection with conventional tools (samtools view -u -M -L configured with required_fields to retain read names, flags, reference identifiers, positions, CIGAR strings and bases, piped into samtools fasta and a k-mer counter given the same query) reduced CPU time 10.5–11.9-fold relative to whole-file streaming. The counting engine reduced CPU time a further 3.9–4.3-fold on the same target regions, with identical counts. Within the engine, single-feature ablations on the region pass resolved three overlapping components: decode-coupled counting in place of inter-process pipes and text formatting (1.9–2.1-fold), the engine’s slice reader and decoder with per-slice reference counting (1.6–1.8-fold), and 16-base Bloom prefiltering with the per-slice reference counting it enables (1.5-fold).

### S7 The Y array

The DYZ3 set comprises 3,816 canonical 31-mers selected across the 41 HPRC training haplotypes. Its k-mers occur at 0.64 copies per assembled base with a coefficient of variation of 0.8% across arrays, so *E* is proportional to length, and in 20 XX genomes the DYZ3 count stayed below 0.14% of the XY median. The calibration factor *q* is the median of *E*/*L* over the 59 training XY genomes (41 HPRC, 18 HGSVC3; family bootstrap 95% interval ±1.1%). In five-fold cross-validation over these genomes, the zero-intercept linear model had the lowest median absolute percentage error of three calibration forms and the linear model with intercept the lowest RMSE; Yruler [12] was run in its region mode on the reads of all 70 XY genomes with assembled arrays (Table S7). Of the 1,259 1000 Genomes genomes recorded as male, 1,255 received an estimate.

### S8 Cloud retrieval and families

In the *All of Us* Researcher Workbench [9], each task copied one DRAGEN-aligned CRAM file to local disk and executed whole-file extraction, targeted extraction on the local copy and targeted extraction by range requests from the storage bucket; summary statistics were aggregated across at least 59 randomly sampled files (Table S12). Because DRAGEN maps a substantial fraction of reads to alternative contigs containing negligible centromeric or normalization k-mers, the DRAGEN-adapted interval list excludes alternative, HLA and EBV contigs.

The trios are the 427 complete parent–offspring trios of the high-coverage 1000 Genomes cohort with counts for all three members. The child’s estimate was regressed on the mid-parent estimate; the slope, corrected for the correlation between spouses, gives *R* (2,000 bootstrap resamples of families). for each trio member, panel estimation excluded training haplotypes from the same 1000 Genomes pedigree and from any training individual with KING kinship *>* 0.05. for the X arrays (per copy), the slope of a daughter’s estimate on her mother’s was 0.43 (0.31–0.56; expected 0.5).

#### Transmission-slope bounds

Each adopted panel target was required to keep the trio transmission slope *R* above that of the corresponding specific-k-mer count minus an target-specific margin from a pedigree simulation on the 427 trios (100 simulated data sets; margin equal to the smaller of −0.05 and the 5th percentile of the simulated 97.5th percentiles of the slope difference; margins range from −0.46 to −0.05). Non-inherited noise lowers *R* below one, and haplotype-correlated bias co-inherited with the array raises it above one; the largest autosomal value is *R* = 1.131 (chr6).

### S9 Population-scale profiling

Of the 3,202 sequenced genomes, six failing sequencing-depth quality control and three discordant with recorded pedigree sex were excluded; among 2,583 pedigree founders, pruning one individual from each of 29 pairs with KING kinship *>* 0.0884 yielded 2,557 unrelated genomes. Variance components were estimated via nested AMOVA on log-transformed array lengths with 1,000 within-population bootstrap resamples; the genome-wide SNV benchmark applied the identical estimator across 114,338 LD-pruned biallelic SNVs (minor allele frequency ≥ 5%, excluding long-range LD regions; Table S14; fig. S3). The residual-bias assessment compares mean out-of-fold training residuals across superpopulations against half the inter-superpopulation median range (using cross-validated interval residuals for chr13, chr21 and family sums) and requires at least two superpopulations with ≥ 5 training genomes, a sample-size requirement met by all targets except chr18 and the chr1+chr5+chr19 family sum. Adjusting each superpopulation median by its mean training residual shifted the difference from the nearest superpopulation from 0.65 to 0.36 Mb (chr21), from 0.81 to 0.61 Mb (chr13 plus chr21), from 1.08 to 1.33 Mb (chr7) and from 0.23 to 0.26 Mb (chr16); every difference kept its direction after the adjustment, and the chr7 difference widened. Depth per copy relative to coverage differed between population medians by at most 0.8%, and females and males within populations by at most 0.016 in log length on any autosomal array. Genome-mean differences between superpopulations appear in assembled lengths in the same direction; estimation residuals also differ between superpopulations (Table S15).

### S10 Association scan in MAGE

MAGE provides RNA sequencing of lymphoblastoid cell lines from 731 donors of the 1000 Genomes Project, with LeafCutter [44] intron-excision ratios grouped by gene, alongside standard covariates (sex, five genotype principal components and PEER factors) [17]; one duplicated library pair (NA11919 and HG00237) was deduplicated to a single donor, leaving 730 donors. Using tensorQTL [45], each array’s length (Blom inverse-normal transformed; chrX normalized per copy within sex) was tested against gene expression and splicing across the array’s chromosome, within 10 Mb of its centromere, and trans-chromosomally (*p <* 5 × 10^−8^), reporting lead gene-level beta-approximated permutation *p* values and Storey *q* values per scan. Detailed results for X-array length and *MED14* splicing are shown in Table S16. Across the 23 tested arrays, 28 cis-chromosomal or pericentromeric associations at *q <* 0.05 (15 expression and 13 splicing; 26 distinct genes) were identified across 11 arrays (Main fig. 6a). Of these, 23 (robust associations) maintained directional consistency across at least four of the five superpopulations (meta-analysis *p <* 0.05) and remained nominal (*p <* 0.05) both without PEER covariates and on untransformed lengths. To evaluate confounding by local linkage disequilibrium, we conditioned each association on the three variants within 1 Mb most strongly associated with the target phenotype. We also constructed a synthetic tag-variant control from the lead SNV plus Gaussian noise. An association met the local-conditioning criterion if it retained at least 50% of its marginal correlation (*p <* 0.05) while the synthetic control retained less than 50% of its marginal correlation. Eleven of the 23 associations met this criterion in the primary scan. for the lead *MED14* intron and the intron shared by *AP5S1* and ENSG00000274949, the initial synthetic controls retained at least 50% of their correlations. We therefore evaluated these two controls across 1,000 independent noise realizations. The *MED14* intron retained 94% of its unadjusted correlation, whereas its synthetic control retained a median of 0.19 and fell below 0.50 in 92% of draws. The ENSG00000274949 control retained a median fraction of 0.20, with less than 0.50 retained in more than 90% of draws.

### S11 Mechanistic basis of Poisson weighting

The reference panel fits normalized k-mer abundances *y_k_* to *μ_k_* = Σ*_a_ β_a_ P_ak_* by weighted non-negative least squares with *w_k_* = max(*μ_k_*, 1)^−1^. Efficient weights are inversely proportional to the conditional variance, Var(*y_k_* | *μ_k_*) = *a μ^γ^* , where *γ* = 1 describes Poisson shot noise and *γ* = 2 quadratic overdispersion. We generated synthetic spectra with variance exponents from 1.0 to 2.0. Using 20 equal-count abundance bins, we estimated these exponents with an absolute error of at most 0.002. We used this binned variance–mean diagnostic on 220 sequenced training genomes (fig. S4a,b).

The panel includes arrays with different HOR subtypes, allowing it to account for differences in k-mer composition that scale with array length. Replacing a single mean-density profile with a panel of assembled arrays reduced the residual variance of k-mer abundances 9.8-fold (median across 16 groups; range 3.3–32.8-fold; fig. S4b).

Weights set how strongly each k-mer influences the length estimate: leverage scales as *w_k_ μ*^2^, proportionally to copy abundance under Poisson weights and uniformly under quadratic weights. High-copy core HOR k-mers track array length (median *r* = 0.95 in the top abundance quintile) and the lowest-abundance quintile correlates weakly with array length (*r* = 0.10; fig. S4c), so Poisson weights concentrate the fit on informative core repeats.

A benchmark of ten weighting schemes on 21 autosomes (five-fold cross-validation; fig. S4d) shows that Poisson weighting combines near-best typical accuracy with robust worst-case accuracy. Poisson weighting has a median *r* of 0.972, a mean *r* of 0.964 and a mean RMSE of 0.261 Mb. The median and mean correlations are within 0.003 of the best scheme for each measure (median *r* 0.974 for unweighted; mean *r* 0.967 for *γ* = 1.25). Its lowest per-autosome correlation (0.865) exceeds that of unweighted (0.747), square-root (0.785), quadratic (0.749), negative-binomial (0.802–0.826) and empirical inverse-variance weighting (0.537); only *γ* = 1.25 (0.890) and the Poisson variant without the lower bound (0.870) reach a higher minimum, and *γ* = 1.25 lowers the correlation significantly on 4 autosomes relative to Poisson weighting. Quadratic weighting lowers mean *r* to 0.951 and raises mean RMSE by 10.4%; it lowers the correlation on 17 of 21 autosomes (11 beyond the paired-bootstrap 95% interval) and raises it on 4 (none beyond the interval). Negative-binomial weighting with *α* = 0.05 lowers the correlation significantly on 8 autosomes and raises it significantly on 1.

Two properties explain the robustness. Empirical inverse-variance weights assign the largest weights to k-mers that are absent from most genomes, whose sample variance is small, and thereby let rare, individual-specific k-mers dominate the fit; the low correlation on the shared-HOR family chr13–chr21 (*r* = 0.537) reflects this. Poisson weights depend on expected abundance only, so a k-mer’s influence follows its copy number in the reference profiles, and the lower bound max(*μ_k_*, 1) limits the weight of k-mers with near-zero expected abundance, which otherwise inflate errors (without this lower bound, chr3 reached an RMSE of 11.8 Mb during model development). When the k-mer sets of different chromosomes do not overlap, Poisson-weighted least squares reduces to the sum of chromosome-specific counts.

### S12 Technical reproducibility across sequencing centres, library preparations and sample composition

The SEQC2 consortium sequenced the breast cancer cell line HCC1395 and the lymphoblastoid cell line HCC1395BL from the same donor at several centres under controlled designs [41, 42]. cenlen read the target regions of all 103 short- read alignments on the NCBI fTP site (BWA-MEM on GRCh38.d1.vd1, with duplicates marked and retained in the files): 42 PCR-free whole-genome libraries, 14 low-input libraries, 21 tumour–normal mixtures and 26 formalin-fixed paraffin-embedded (ffPE) libraries. The alignment build lacks the 786 alternative and HLA target intervals; in one library read in full, estimates changed by a median 0.13%. The genome-wide scale *g* of a library is the median over autosomes of log(*L*^^^/*L*), where *L* is the median estimate across the cell line’s PCR-free libraries.

#### Sequencing centres and platforms

Each cell line has 22 PCR-free libraries from six centres on HiSeq X Ten, HiSeq 4000 and NovaSeq 6000 instruments. for relative lengths (Methods), the replicate SD of log length had a median of 0.021 in HCC1395BL and 0.020 in HCC1395 across autosomes (fig. 3e), against 0.21 across the 2,557 unrelated 1000 Genomes genomes; reliability, 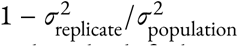, had a median of 0.99 in both cell lines. The two lines come from one donor, and their relative lengths differ beyond the replicate ranges on 18 of 22 autosomes, consistent with copy-number changes in the tumour line. for absolute lengths, the median reliability was 0.91 and 0.93: one HiSeq 4000 batch of three libraries per cell line had *g* of +0.055 to +0.187, whereas every other PCR-free library, including the NovaSeq libraries of the same centre, had | *g* | *<* 0.05 (fig. S6a). Without this batch, the median reliability of absolute lengths was 0.99, and the leave-one-out consensus lay within the 95% prediction interval in 97.8% of 874 library–array pairs. In the batch, array k-mer occurrences per read matched those of the other centres and single-copy k-mer occurrences per read were lower (fig. S6a), so the offset arises in the depth normalization and is removed in relative lengths.

#### Library preparation

Low-input libraries were prepared with TruSeq-Nano (1, 10 and 100 ng) and TruSeq PCR- free (250 ng) at one centre and with Nextera flex (1, 10 and 100 ng) at another. With duplicate-flagged reads retained, as in calibration, TruSeq-Nano libraries were within −0.006 to +0.011 in *g* of the same centre’s PCR-free libraries at duplicate shares up to 0.65 (fig. S6b). Removing duplicate-flagged reads raised *g* in every low-input library, to as much as +1.50 at 1 ng. The increase occurred because duplicate marking removed a smaller share of array k-mer occurrences than of single-copy occurrences. We observed this pattern in 98 of the 103 libraries. Nextera flex libraries had *g* of +0.02 to +0.08; all six come from one centre, so kit and centre effects coincide for this kit.

#### FFPE

The 26 ffPE libraries (fixation for 1, 2, 6 and 24 h at two centres) had *g* of −1.43 to +0.77 and array-specific residual SDs of 0.04 to 0.14 (fig. S6c); cenlen’s calibration applies to DNA from blood, cell lines and fresh or frozen tissue.

#### Tumour–normal mixtures

Mixtures of HCC1395 and HCC1395BL DNA at tumour fractions *f* of 0.05, 0.10, 0.20, 0.50 and 0.75 (three libraries each) were compared with a prediction from the six unmixed libraries alone: the counts per read of each query k-mer were mixed as *f u_T_* + (1 − *f*)*u_N_* and passed through cenlen, which for an estimator linear in the counts gives *L*_mix_(*f*) = [ *f d_T_ L_T_* + (1 − *f*) *d_N_ L_N_* ]/[ *f d_T_* + (1 − *f*) *d_N_* ], where *d* is the single-copy depth per read (*d_T_* /*d_N_* = 1.154). After removal of each library’s scale *g*, the median root-mean-square error of log length across arrays was 0.0067 for this prediction and 0.0103 for linear mixing of the two lengths, close to the replicate SD of the unmixed libraries (0.0058); the prediction was closer on 20 of 22 autosomes (two-sided sign test, *p* = 1.2 × 10^−4^; fig. S6d). With the scale retained, the errors were 0.0136 and 0.0158.

---

**Figure S1:**
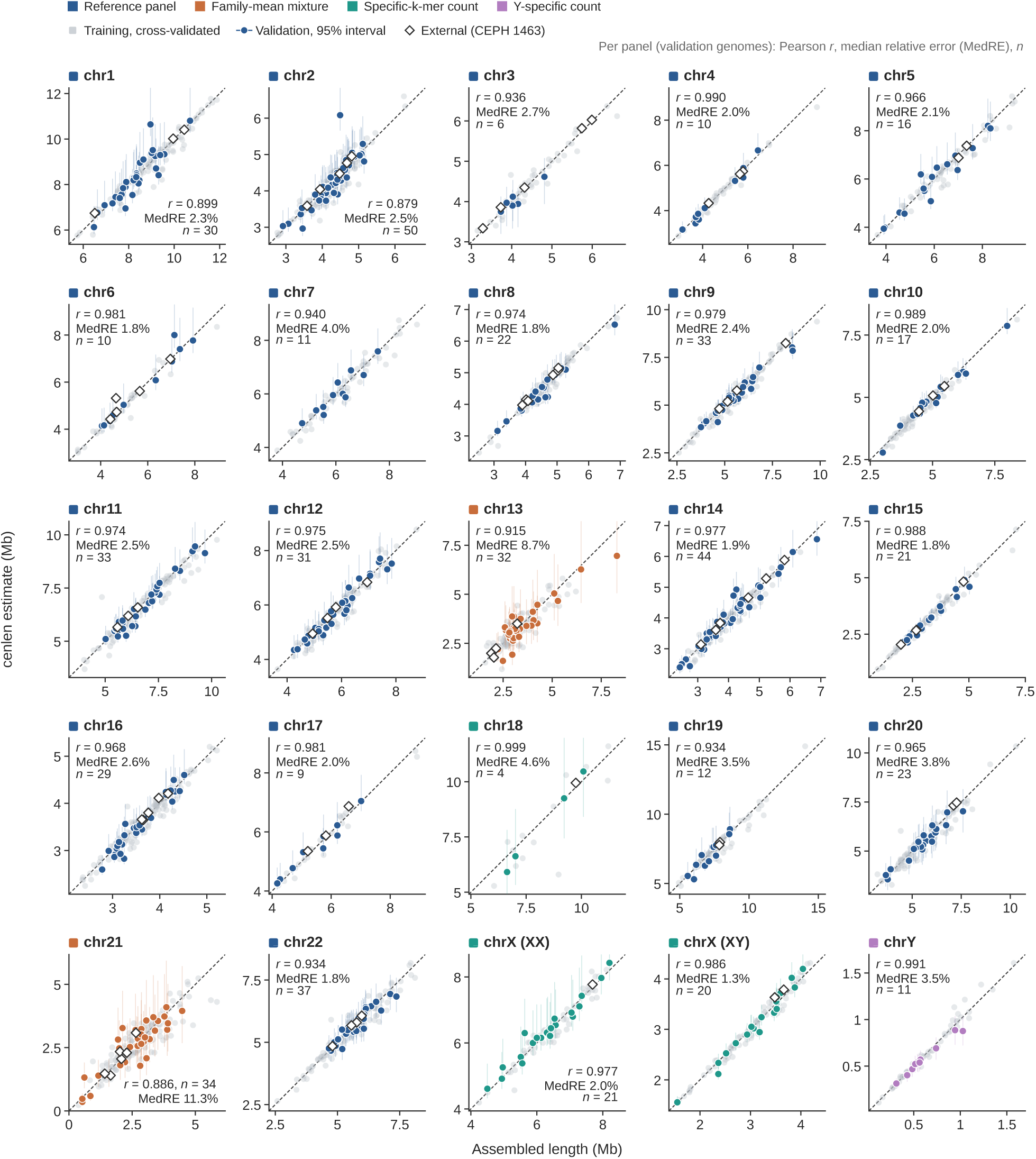
Estimates against assembled length for all 25 targets. Each panel shows cenlen estimates against assembled length for training genomes in five-fold cross-validation across pedigree families (grey; 219 genomes, and for chrY the 59 training XY genomes of the Y-array calibration), validation genomes with 95% prediction intervals (61 genomes, 566 genome–array pairs; colours mark the model, Table S4) and CEPH 1463 genomes (open diamonds; 6 genomes, 83 pairs), with the identity line (dashed). Each panel gives the Pearson correlation, median relative error (MedRE) and number of validation genomes. Autosomes and the X chromosome of XX genomes are sums of two arrays; the X chromosome of XY genomes and the Y array are single copies.

**Figure S2:**
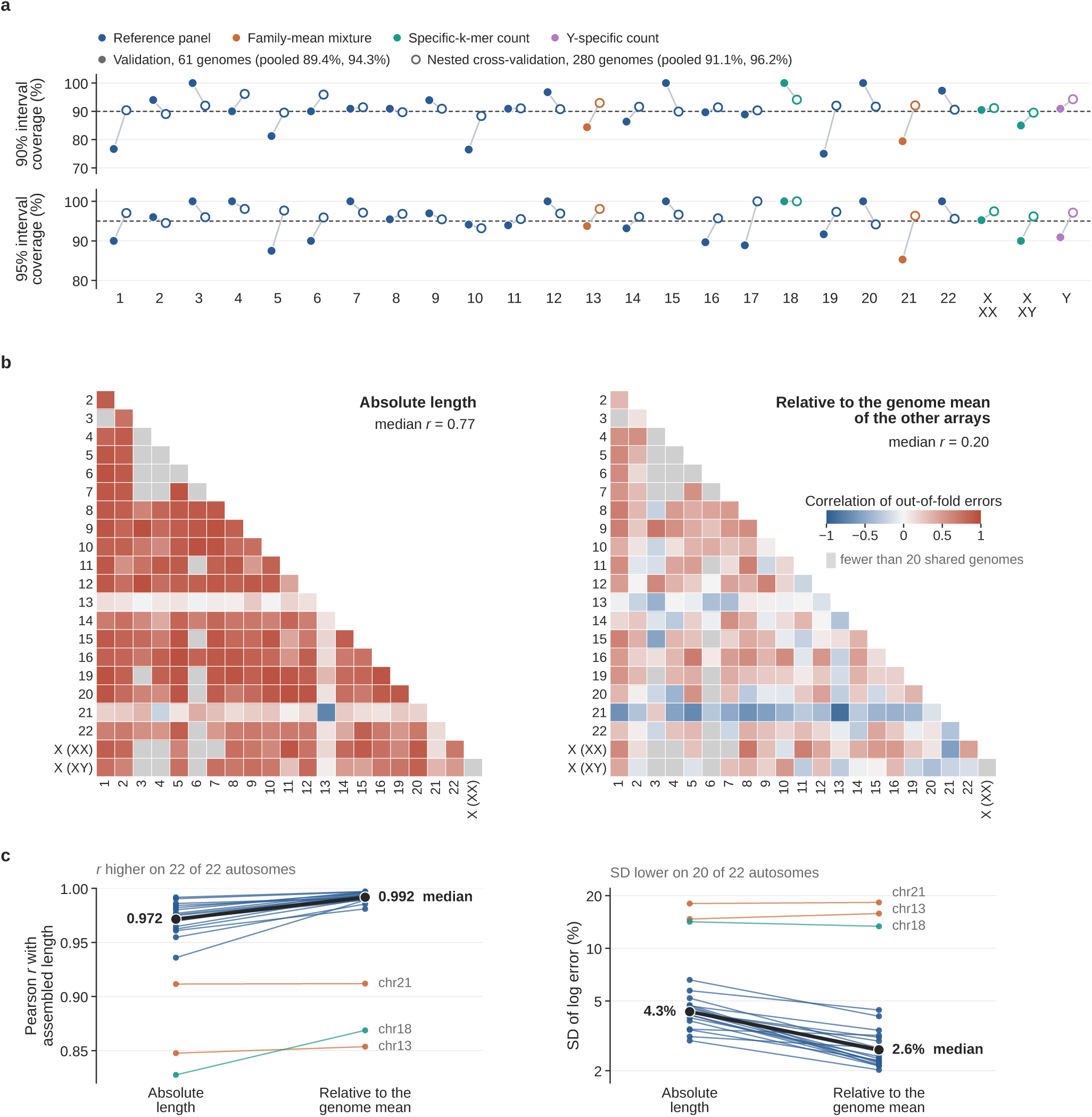
Prediction-interval coverage and the genome-level error component shared across arrays. (a) Empirical coverage of the 90% and 95% prediction intervals per target on the 61 validation genomes (filled; pooled 89.4% and 94.3%, 95% family-cluster bootstrap intervals 84.7–93.2% and 91.5–96.7%) and in five-fold nested cross-validation across 280 sequenced genomes (open; pooled 91.1% and 96.2%), coloured by model; dashed lines mark nominal levels. (b) Correlation across 219 training genomes between the out-of-fold relative errors of each pair of targets for absolute lengths (left; median *r* = 0.77) and for lengths relative to the genome mean of the other arrays (right; median *r* = 0.20); grey, pairs with fewer than 20 shared genomes; chr17, chr18 and chrY, which share fewer than 20 genomes with every other target, are omitted. (c) Correlation with assembled length and standard deviation of the log error per autosome for absolute lengths and for lengths relative to the genome mean of the other autosomes (black, medians: 0.972 to 0.992; 4.3% to 2.6%) in 192 training genomes with at least six assembled autosomes; the correlation is higher on all 22 autosomes and the SD lower on 20.

**Figure S3:**
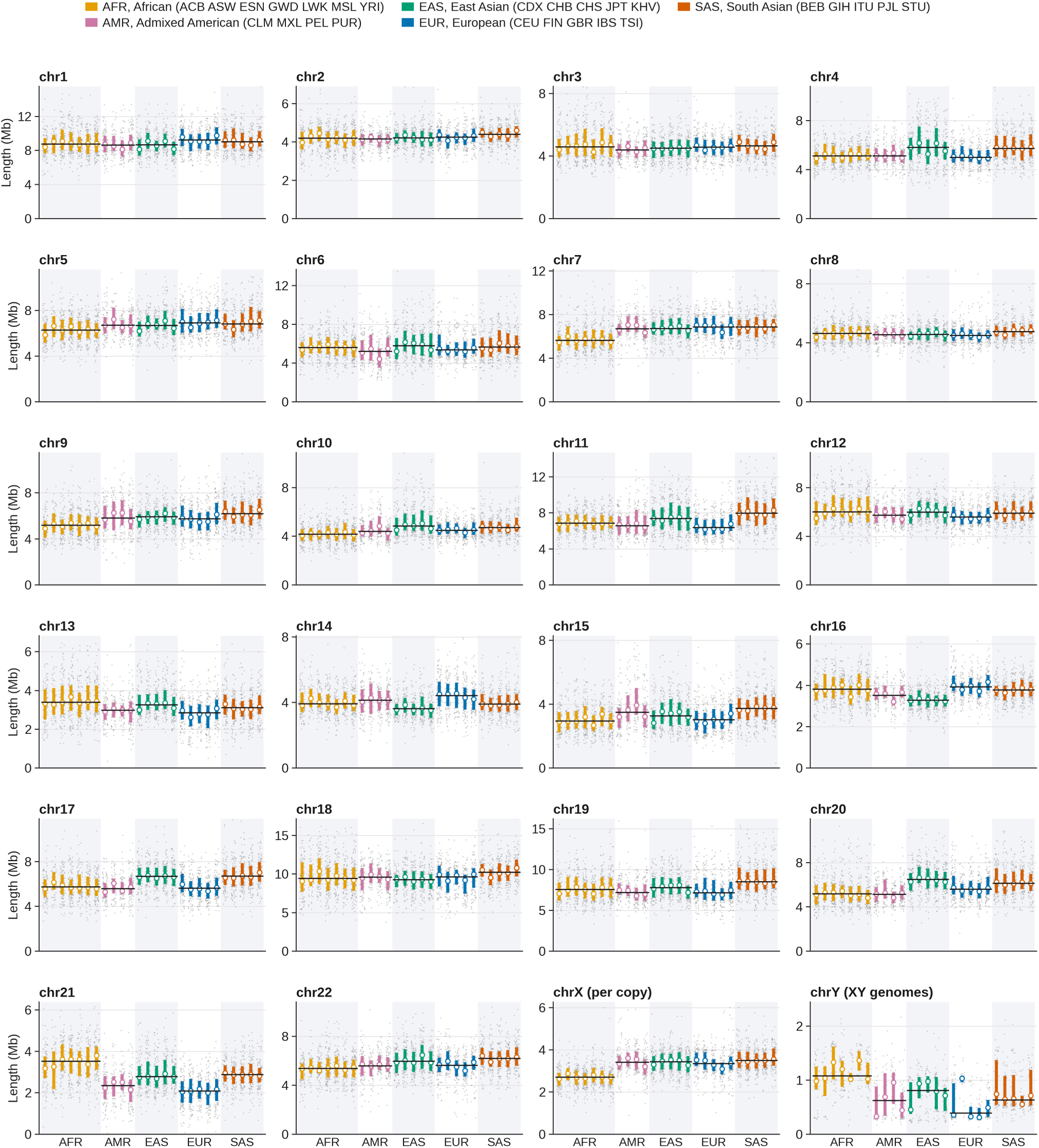
Array length of every chromosome in the 26 populations of the 1000 Genomes Project. cenlen estimates in 2,557 unrelated genomes: diploid totals for chr1–chr22, the X array per copy (XX totals halved; both sexes) and the Y array of 1,268 XY genomes. Grey points denote genomes; for each population, the coloured bar spans the interquartile range and the open dot marks the median; black lines mark superpopulation medians. Populations are ordered within superpopulation as listed in the key (AfR, African; AMR, admixed American; EAS, East Asian; EUR, European; SAS, South Asian), the order of fig. 5b. Vertical axes start at zero and end at 1.06 times the 99.8th percentile of each panel (at most five genomes per panel lie above).

**Figure S4:**
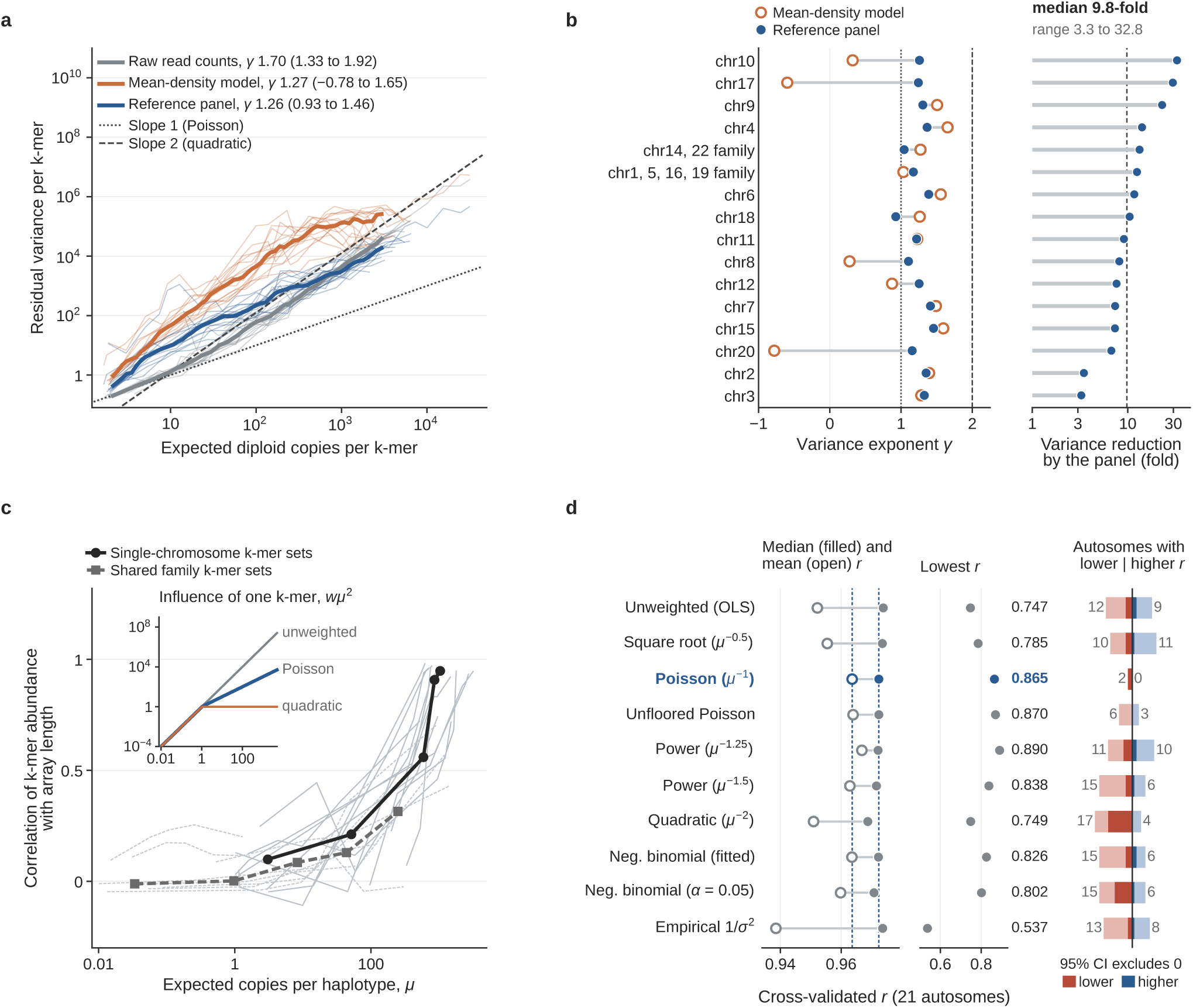
Variance structure of k-mer abundances and the choice of Poisson weights. (a) Residual variance of normalized k-mer abundances against expected diploid copies in 20 equal-count bins for each of 16 k-mer groups (thin lines; thick, median across groups), for raw read counts around each genome’s own assembly, the cross-validated mean-density model and reference panels (220 sequenced training genomes); dotted and dashed lines, slopes 1 (Poisson) and 2 (quadratic); *γ*, median (range) of the fitted variance exponent across groups. (b) fitted variance exponent per k-mer group under the mean-density model and reference panels (left), and the fold reduction of summed residual variance by the panel (right). (c) Correlation of k-mer abundance with array length by abundance quintile of each array’s k-mers (thin lines; thick, medians) for single-chromosome and shared family k-mer sets; inset, influence of one k-mer on the fit, *wμ*^2^, under unweighted, Poisson and quadratic weights. (d) Ten weighting schemes (*w_k_* = max(*μ_k_*, 1)^−^*^γ^* for power schemes) in five-fold cross-validation across 21 autosomes: median and mean correlation with assembled length (dashed, Poisson), lowest correlation, and the number of autosomes with lower or higher correlation than the released weights (Poisson; without the lower bound for the chr13–chr21 family-mean mixture), darker where the paired-bootstrap 95% interval of the difference excludes zero.

**Figure S5:**
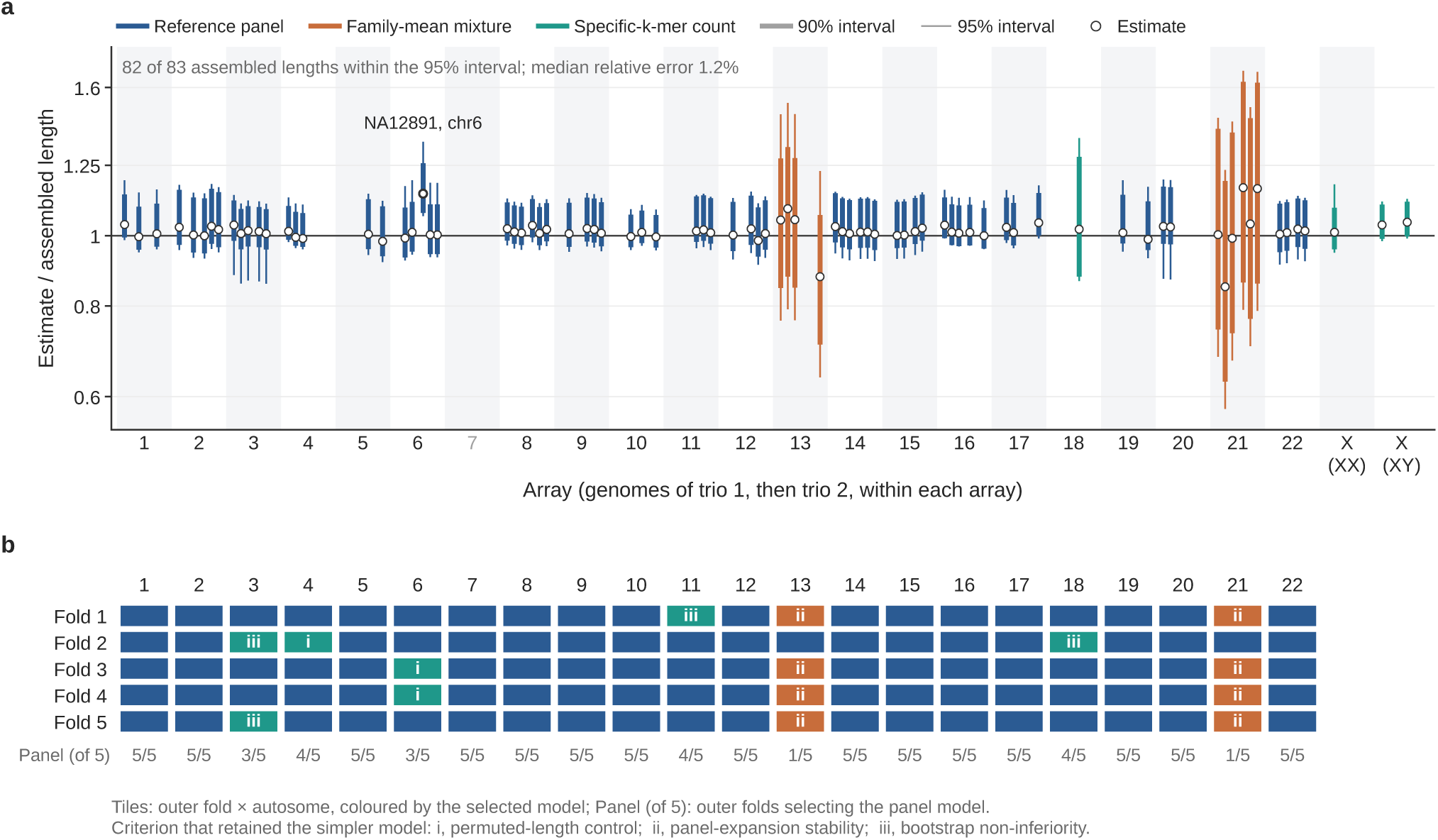
Validation in CEPH 1463 genomes and model assignment in nested cross-validation. (a) cenlen estimates relative to Platinum Pedigree assembled lengths, with 90% (thick) and 95% (thin) prediction intervals, for the six CEPH 1463 genomes and every genome–array pair with a completely assembled reference (83 pairs; none for chr7 and chrY). Genome order within each array: NA12889, NA12890, NA12877 (trio 1) and NA12891, NA12892, NA12878 (trio 2); colours mark the model; the single assembled length outside its 95% interval is labelled. (b) Model assigned to each autosome by the automated rule in each outer fold of five-fold family-stratified nested cross-validation (282 sequenced genomes), aligned with (a); for folds where the simpler model was selected, the label identifies the criterion that led to its selection (Supplementary Section S5): i, permuted-length control; ii, stability under panel expansion with assembly-only haplotypes; iii, bootstrap non-inferiority. The released assignment retains chr18 at the specific-k-mer count by a small-sample rule.

**Figure S6:**
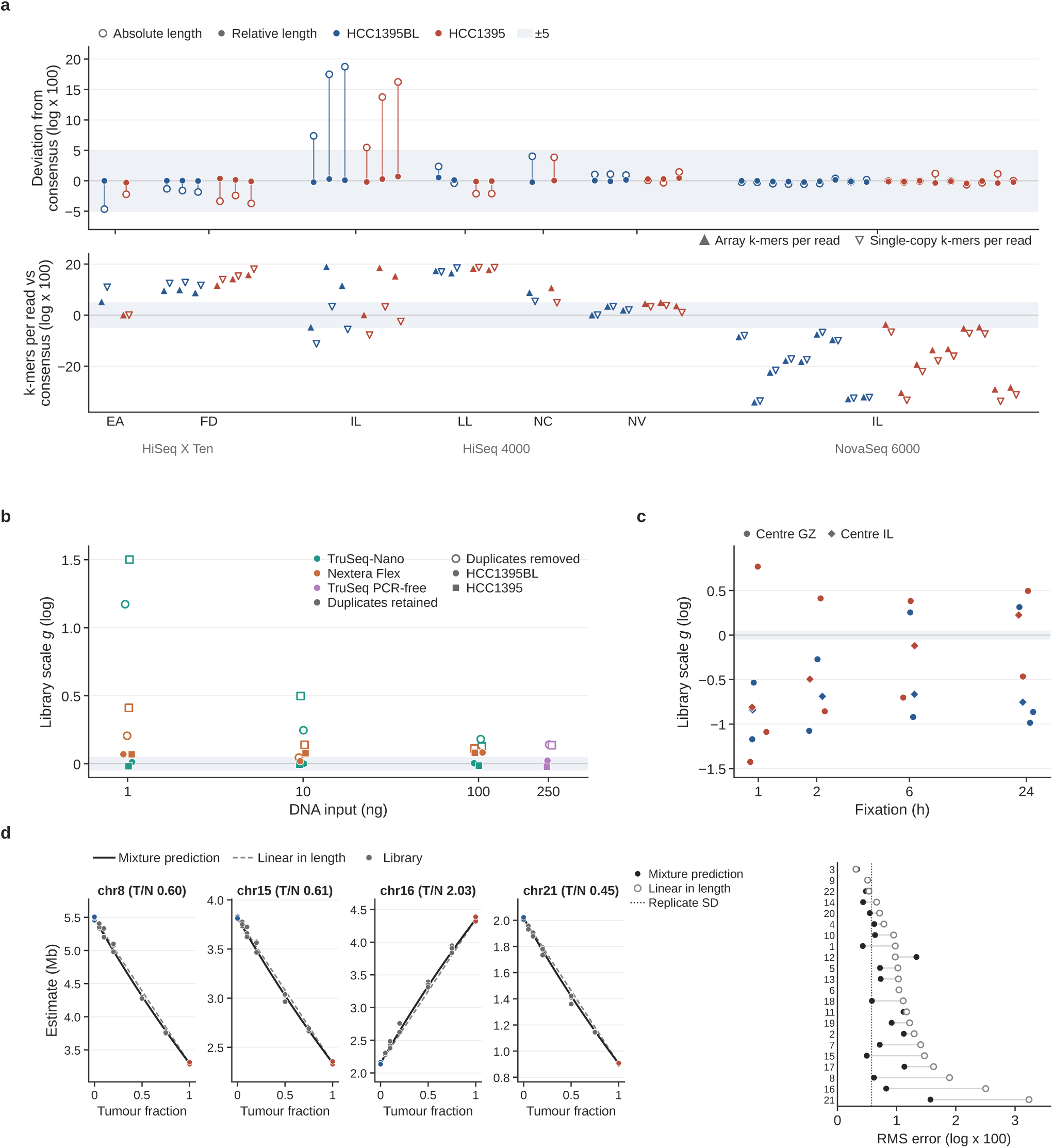
Technical reproducibility across 103 SEQC2 libraries of HCC1395 and HCC1395BL. (a) Top, deviation of each PCR-free library from its cell line’s consensus for absolute length (open) and relative length (filled), median over autosomes, joined per library and grouped by sequencing centre and platform (IL under NovaSeq 6000 marks the NovaSeq libraries of the centre whose HiSeq 4000 batch reads high). Bottom, in the same library order, array and single-copy k-mer occurrences per read relative to the median of the other centres’ PCR-free libraries; both rows in log units ×100, shaded ±5. (b) Genome-wide scale *g* of low-input libraries by DNA input and kit, with duplicate-flagged reads retained (filled) and removed (open); circles, HCC1395BL; squares, HCC1395. (c) *g* of ffPE libraries by fixation time; circles, centre GZ; diamonds, centre IL; blue, HCC1395BL; red, HCC1395. (d) Tumour–normal mixtures: estimates along tumour DNA fraction for the four autosomal arrays with the largest tumour/normal contrast (T/N, ratio of the unmixed means), with the mixture prediction from the unmixed libraries’ counts per read (solid) and linear mixing of the two lengths (dashed); right, root-mean-square error of log length for both predictions on all 22 autosomes after removal of each library’s scale, with the replicate SD of the unmixed libraries (dotted) (Supplementary Section S12).

**Figure S7:**
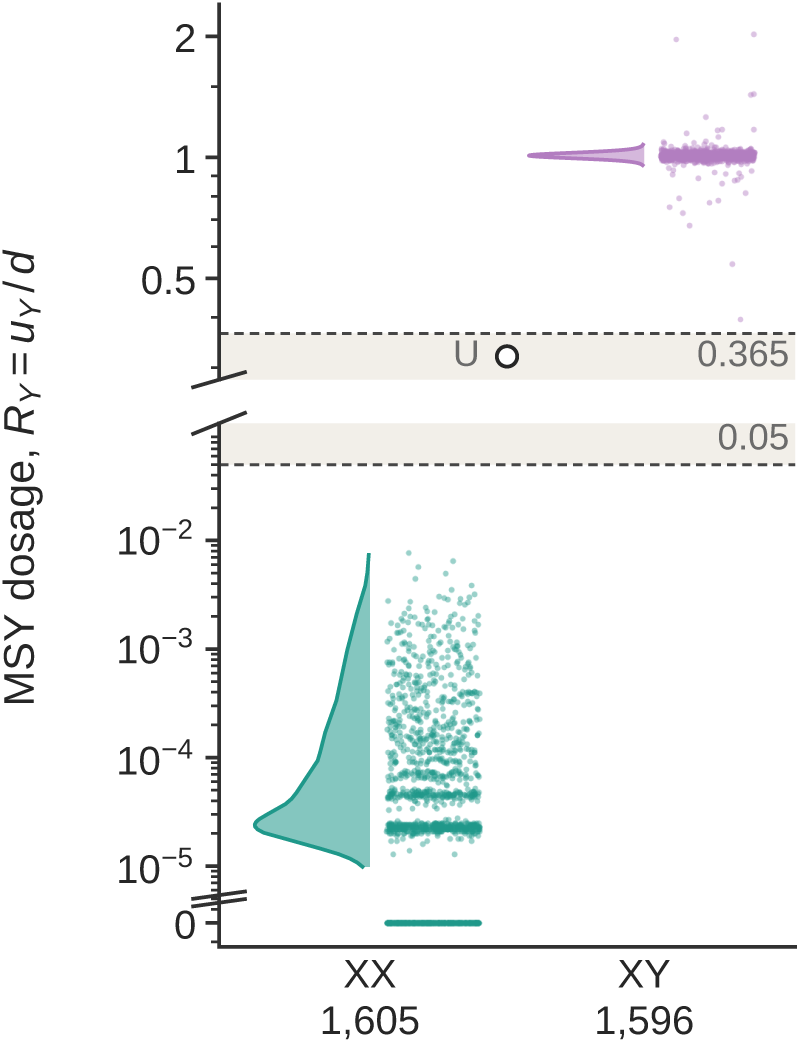
MSY dosage by inferred sex. MSY dosage *R_Y_* = *u_Y_* /*d* in 3,202 high-coverage 1000 Genomes genomes (U, unclassified; XX range 0.0000–0.0078, XY range 0.3960–2.0299): half-violins, density of log *R_Y_* ; points, genomes; log axis broken between 0.12 and 0.28; dashed lines, thresholds 0.05 and 0.365; 503 XX genomes without MSY k-mer counts at 0.

**Table S1:** Accuracy per target. Training genomes: five-fold cross-validation across pedigree families. Validation genomes: the 61 validation genomes with complete reference arrays (five-fold nested cross-validation across all 282 sequenced genomes is reported in Table S2). Model: Panel, reference panel; Mixture, family-mean mixture; Set, specific-k-mer count; Y ratio, Y-specific count (DYZ3 count normalized by single- copy male-specific chrY k-mers). MedRE, median absolute relative error; RMSE, root-mean-square error; 95% cov., share of assembled lengths within the 95% prediction interval; *R*, Mendelian transmission slope across 427 trios.

|  |  | Training, cross-validated |  |  | Validation |  |  |  |  | Trios |
|---|---|---|---|---|---|---|---|---|---|---|
| Output | Model | *n* | *r* | MedRE (%) | *n* | *r* | MedRE (%) | RMSE (Mb) | 95% cov. | *R* |
| chr1 | Panel | 105 | 0.970 | 1.8 | 30 | 0.899 | 2.3 | 0.47 | 0.90 | 0.94 |
| chr2 | Panel | 169 | 0.934 | 2.7 | 50 | 0.879 | 2.5 | 0.29 | 0.96 | 0.90 |
| chr3 | Panel | 44 | 0.965 | 2.7 | 6 | 0.936 | 2.7 | 0.14 | 1.00 | 1.01 |
| chr4 | Panel | 42 | 0.991 | 1.4 | 10 | 0.990 | 1.9 | 0.17 | 1.00 | 0.85 |
| chr5 | Panel | 70 | 0.974 | 1.8 | 16 | 0.966 | 2.1 | 0.34 | 0.88 | 0.98 |
| chr6 | Panel | 39 | 0.977 | 2.2 | 10 | 0.981 | 1.8 | 0.30 | 0.90 | 1.13 |
| chr7 | Panel | 59 | 0.976 | 2.2 | 11 | 0.940 | 4.0 | 0.27 | 1.00 | 0.91 |
| chr8 | Panel | 104 | 0.975 | 1.7 | 22 | 0.974 | 1.8 | 0.18 | 0.95 | 0.94 |
| chr9 | Panel | 121 | 0.983 | 2.3 | 33 | 0.979 | 2.4 | 0.25 | 0.97 | 0.91 |
| chr10 | Panel | 86 | 0.986 | 1.4 | 17 | 0.989 | 2.0 | 0.18 | 0.94 | 0.95 |
| chr11 | Panel | 101 | 0.963 | 1.8 | 33 | 0.974 | 2.5 | 0.30 | 0.94 | 1.02 |
| chr12 | Panel | 99 | 0.968 | 2.5 | 31 | 0.975 | 2.5 | 0.23 | 1.00 | 0.88 |
| chr13 | Mixture | 124 | 0.872 | 8.2 | 32 | 0.915 | 8.7 | 0.51 | 0.94 | 0.99 |
| chr14 | Panel | 135 | 0.974 | 2.3 | 44 | 0.977 | 1.9 | 0.20 | 0.93 | 1.00 |
| chr15 | Panel | 98 | 0.990 | 1.7 | 21 | 0.988 | 1.8 | 0.16 | 1.00 | 0.96 |
| chr16 | Panel | 134 | 0.972 | 2.0 | 29 | 0.968 | 2.6 | 0.15 | 0.90 | 0.89 |
| chr17 | Panel | 22 | 0.987 | 1.7 | 9 | 0.981 | 2.0 | 0.18 | 0.89 | 0.95 |
| chr18 | Set | 13 | 0.852 | 9.3 | 4 | 0.999 | 4.6 | 0.45 | 1.00 | 0.96 |
| chr19 | Panel | 63 | 0.972 | 1.8 | 12 | 0.933 | 3.5 | 0.40 | 0.92 | 1.00 |
| chr20 | Panel | 97 | 0.960 | 2.7 | 23 | 0.965 | 3.8 | 0.27 | 1.00 | 0.99 |
| chr21 | Mixture | 130 | 0.891 | 10.5 | 34 | 0.886 | 11.3 | 0.47 | 0.85 | 0.96 |
| chr22 | Panel | 122 | 0.968 | 2.4 | 37 | 0.933 | 1.8 | 0.22 | 1.00 | 0.93 |
| chrX (XX) | Set | 58 | 0.972 | 1.7 | 21 | 0.977 | 2.0 | 0.21 | 0.95 | – |
| chrX (XY) | Set | 85 | 0.987 | 1.9 | 20 | 0.986 | 1.3 | 0.11 | 0.90 | – |
| chrY | Y ratio | 59 | 0.990 | 2.5 | 11 | 0.991 | 3.5 | 0.05 | 0.91 | – |

**Table S2:** Five-fold family-stratified nested cross-validation across all 282 sequenced genomes. Model selection, calibration and prediction intervals were refitted independently within each outer training split (280 genomes with complete reference arrays). Panel folds: number of outer folds in which the automated selection rule assigned the reference panel. Unrelated to k-mer selection: held-out genomes whose pedigree and kinship units contain none of the genomes used to select k-mers. Bottom rows report shared-HOR family totals.

|  |  |  | All held-out genomes (*n* = 280) |  |  |  | Unrelated to k-mer selection |  |
|---|---|---|---|---|---|---|---|---|
| Output | *n* | Panel folds | *r* | MedRE (%) | RMSE (Mb) | 95% cov. | MedRE (%) | *n* |
| chr1 | 135 | 5/5 | 0.959 | 1.9 | 0.37 | 0.97 | 2.0 | 77 |
| chr2 | 219 | 5/5 | 0.921 | 2.8 | 0.23 | 0.95 | 2.7 | 130 |
| chr3 | 50 | 3/5 | 0.960 | 2.6 | 0.20 | 0.96 | 2.4 | 25 |
| chr4 | 52 | 4/5 | 0.983 | 2.1 | 0.21 | 0.98 | 2.2 | 27 |
| chr5 | 86 | 5/5 | 0.973 | 1.8 | 0.30 | 0.98 | 1.8 | 47 |
| chr6 | 49 | 3/5 | 0.979 | 2.3 | 0.27 | 0.96 | 1.8 | 26 |
| chr7 | 70 | 5/5 | 0.972 | 2.3 | 0.27 | 0.97 | 2.7 | 34 |
| chr8 | 126 | 5/5 | 0.974 | 1.7 | 0.16 | 0.97 | 1.8 | 78 |
| chr9 | 154 | 5/5 | 0.981 | 2.4 | 0.24 | 0.95 | 2.5 | 93 |
| chr10 | 103 | 5/5 | 0.987 | 1.6 | 0.15 | 0.93 | 1.5 | 54 |
| chr11 | 134 | 4/5 | 0.965 | 2.0 | 0.35 | 0.96 | 2.1 | 82 |
| chr12 | 130 | 5/5 | 0.970 | 2.4 | 0.24 | 0.97 | 2.5 | 78 |
| chr13 | 156 | 1/5 | 0.907 | 7.3 | 0.42 | 0.98 | 7.6 | 94 |
| chr14 | 179 | 5/5 | 0.976 | 2.3 | 0.17 | 0.96 | 2.0 | 104 |
| chr15 | 119 | 5/5 | 0.989 | 1.8 | 0.16 | 0.97 | 2.0 | 70 |
| chr16 | 163 | 5/5 | 0.971 | 2.1 | 0.14 | 0.96 | 2.2 | 90 |
| chr17 | 31 | 5/5 | 0.986 | 1.7 | 0.20 | 1.00 | 1.6 | 15 |
| chr18 | 17 | 4/5 | 0.950 | 3.7 | 0.57 | 1.00 | 3.2 | 6 |
| chr19 | 75 | 5/5 | 0.971 | 1.8 | 0.40 | 0.97 | 2.1 | 40 |
| chr20 | 120 | 5/5 | 0.959 | 3.2 | 0.34 | 0.94 | 3.4 | 66 |
| chr21 | 164 | 1/5 | 0.915 | 8.3 | 0.41 | 0.96 | 8.5 | 109 |
| chr22 | 159 | 5/5 | 0.966 | 2.1 | 0.26 | 0.96 | 1.8 | 88 |
| chrX (XX) | 79 | – | 0.972 | 1.7 | 0.23 | 0.97 | 1.8 | 59 |
| chrX (XY) | 105 | – | 0.987 | 1.9 | 0.10 | 0.96 | 1.9 | 58 |
| chrY | 70 | – | 0.987 | 2.9 | 0.05 | 0.97 | 3.0 | 37 |
| chr13 + chr21 | 93 | – | 0.978 | 2.7 | 0.35 | – | 3.1 | 55 |
| chr14 + chr22 | 106 | – | 0.954 | 1.9 | 0.39 | – | 1.6 | 56 |
| chr1 + chr5 + chr19 | 19 | – | 0.843 | 1.4 | 1.32 | – | 0.6 | 6 |

**Table S3:** Selection and evaluation criteria for chromosome-specific k-mer sets. *μ_k_*, mean copies of k-mer *k* per selection haplotype within the active array; *λ*, stringency factor. Parameter pairs (*p*, *λ*) were evaluated sequentially in the order (0.90, 1), (0.75, 1), (0.50, 1), (0.75, 2) and (0.50, 2), selecting the first configuration yielding ≥ 1,000 k-mers and five-fold cross-validated correlation *r* ≥ 0.80 between summed k-mer copies and assembled array length.

| Criterion | Threshold |
|---|---|
| *Selection (each candidate 31-mer of the active array)* |  |
| Other chromosomes’ centromeres | mean copies ≤ 0.005*λμ<sub>k</sub>* per chromosome and ≤ 0.01*λμ<sub>k</sub>* summed; prevalence ≤ 5*λ*%; 99th percentile ≤ 0.05*μ<sub>k</sub>* |
| Own centromere outside the array | mean copies ≤ 0.01*λμ<sub>k</sub>*; also in T2T-CHM13 outside the active-array intervals |
| Single-substitution neighbours | summed copies of the 93 neighbours ≤ 10*μ<sub>k</sub>* (selection centromeres and T2T-CHM13) |
| Prevalence | ≥ *p* of selection haplotypes; ≥ *p* − 0.10 in each ancestry group with ≥ 10 haplotypes |
| *Post-selection verification (137 non-selection training genomes unrelated to selection set)* |  |
| Copies inside active arrays | ≥ 95% (observed 99.8–100.0%) |
| Occurrences on other chromosomes | ≤ 2% of its own (observed ≤ 0.0011%) |
| Assembled copy number versus array length | *r* ≥ 0.80 per haplotype (observed 0.910–0.999) |
| Short-read counts versus assembly-predicted copies | *r* ≥ 0.85 (observed 0.936–0.999) |

**Table S4:** Model specification for each cenlen target. Listed are the model (as in Table S1), the k-mer set and its size, the number of training arrays in the panel (excluding arrays below half the chromosome-specific median k-mer density) and the calibration factor *f_c_* . for family-panel targets, per-chromosome array counts are listed individually while arrays from all family members enter the joint fit. The chrY model is described in Supplementary Section S7.

| Output | Model | k-mer set | k-mers | Panel arrays | Calibration factor *f<sub>c</sub>* |
|---|---|---|---|---|---|
| chr1 | Panel | family: chr1, chr5, chr16, chr19 | 10,000 | 321 | 0.961 |
| chr2 | Panel | chromosome-specific | 2,097 | 435 | 0.993 |
| chr3 | Panel | chromosome-specific | 3,949 | 185 | 0.987 |
| chr4 | Panel | chromosome-specific | 2,023 | 197 | 1.000 |
| chr5 | Panel | family: chr1, chr5, chr16, chr19 | 10,000 | 253 | 0.980 |
| chr6 | Panel | chromosome-specific | 3,331 | 204 | 0.998 |
| chr7 | Panel | chromosome-specific | 1,029 | 205 | 0.958 |
| chr8 | Panel | chromosome-specific | 1,638 | 328 | 1.017 |
| chr9 | Panel | chromosome-specific | 1,170 | 350 | 0.978 |
| chr10 | Panel | chromosome-specific | 1,025 | 300 | 1.012 |
| chr11 | Panel | chromosome-specific | 1,186 | 307 | 1.017 |
| chr12 | Panel | chromosome-specific | 1,587 | 311 | 0.974 |
| chr13 | Mixture | family: chr13, chr21 | 143,094 | – | 1.019 |
| chr14 | Panel | family: chr14, chr22 | 10,000 | 379 | 1.011 |
| chr15 | Panel | chromosome-specific | 2,236 | 307 | 0.992 |
| chr16 | Panel | family: chr1, chr5, chr16, chr19 | 10,000 | 371 | 0.976 |
| chr17 | Panel | chromosome-specific | 1,741 | 156 | 0.997 |
| chr18 | Set | chromosome-specific | 1,479 | – | 1.017 |
| chr19 | Panel | family: chr1, chr5, chr16, chr19 | 10,000 | 229 | 0.936 |
| chr20 | Panel | chromosome-specific | 1,151 | 290 | 0.977 |
| chr21 | Mixture | family: chr13, chr21 | 143,094 | – | 1.090 |
| chr22 | Panel | family: chr14, chr22 | 10,000 | 358 | 1.007 |
| chrX (XX) | Set | chromosome-specific | 3,181 | – | 0.965 |
| chrX (XY) | Set | chromosome-specific | 3,181 | – | 0.963 |

**Table S5:**
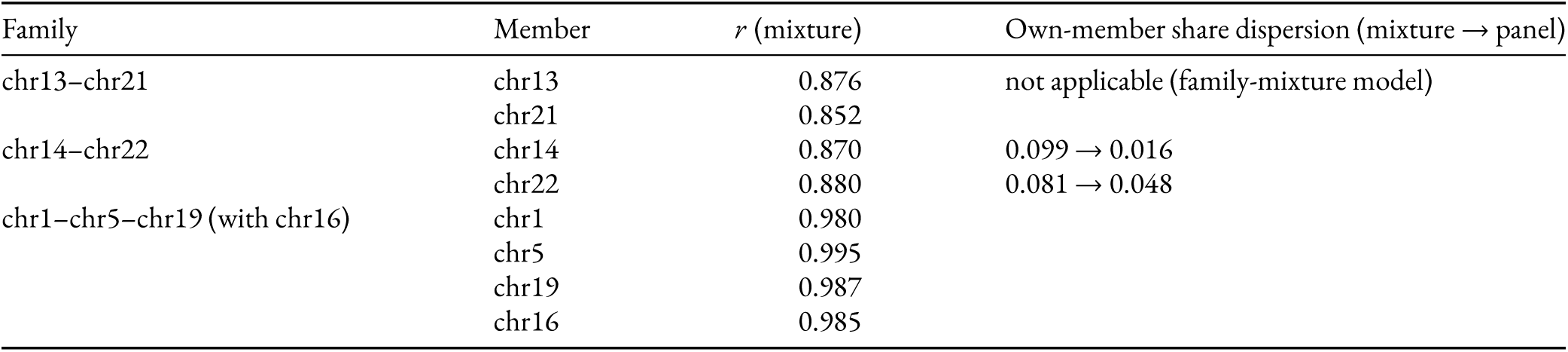
Member separation in synthetic diploid assemblies. Across 400 synthetic diploids per fold constructed from held-out training haplotypes: correlation between family-mean mixture estimates and true member lengths (median relative errors 1.0–7.0%; permuting member labels within chr13–chr21 yielded *r* = 0.29 and 0.10). Right column: dispersion in the fraction of each array’s length assigned to its true chromosome under the family-mean mixture versus the family panel (cross-member misassignment was at most 0.3%).

| Family | Member | *r* (mixture) | Own-member share dispersion (mixture → panel) |
|---|---|---|---|
| chr13–chr21 | chr13 | 0.876 | not applicable (family-mixture model) |
|  | chr21 | 0.852 |  |
| chr14–chr22 | chr14 | 0.870 | 0.099 → 0.016 |
|  | chr22 | 0.880 | 0.081 → 0.048 |
| chr1–chr5–chr19 (with chr16) | chr1 | 0.980 |  |
|  | chr5 | 0.995 |  |
|  | chr19 | 0.987 |  |
|  | chr16 | 0.985 |  |

**Table S6:** Concordance of cenlen and HOROSCOPE with Platinum Pedigree assemblies in six CEPH 1463 individuals. (a) Matched genome–array pairs across the two parent–offspring trios (lengths in Mb; bias, estimate minus assembled length; MedAE (arrays), mean across chromosomes of per-chromosome median absolute errors; MedAE (pairs), median over all pairs). (b) Array-averaged MedAE after leaving out one genome or one whole trio (trio A, NA12877 with his parents NA12889 and NA12890; trio B, NA12878 with her parents NA12891 and NA12892); recalibrated HOROSCOPE fits assembled length on twice the reported HOR length per chromosome across 59 HOROSCOPE training genomes.

| **a** All genome–array pairs (lengths in Mb) |  |  |  |  |  |  |  |
|---|---|---|---|---|---|---|---|
| Method | MedAE (arrays) | MedAE (pairs) | MAE | RMSE | Bias | MedRE (%) |  |
| *Chromosomes with training recalibration, 47 genome–array pairs* |  |  |  |  |  |  |  |
| cenlen | 0.073 | 0.053 | 0.081 | 0.128 | +0.066 | 1.1 |  |
| HOROSCOPE, as published | 0.241 | 0.181 | 0.228 | 0.285 | -0.028 | 3.5 |  |
| HOROSCOPE, recalibrated | 0.251 | 0.160 | 0.212 | 0.298 | +0.065 | 3.6 |  |
| *All 17 HOROSCOPE autosomes, 56 genome–array pairs* |  |  |  |  |  |  |  |
| cenlen | 0.085 | 0.059 | 0.089 | 0.133 | +0.070 | 1.1 |  |
| HOROSCOPE, as published | 0.237 | 0.183 | 0.230 | 0.283 | -0.013 | 3.4 |  |
| **b** Array-averaged MedAE (Mb) after leaving out one genome or one trio |  |  |  |  |  |  |  |
|  | Arrays with HOROSCOPE recalibration |  |  |  | All 17 HOROSCOPE autosomes |  |  |
| Genomes | Pairs | cenlen | HOROSCOPE | Recalibrated | Pairs | cenlen | HOROSCOPE |
| All six genomes | 47 | 0.073 | 0.241 | 0.251 | 56 | 0.085 | 0.237 |
| Without NA12877 | 39 | 0.076 | 0.248 | 0.260 | 46 | 0.096 | 0.244 |
| Without NA12878 | 38 | 0.076 | 0.240 | 0.244 | 45 | 0.082 | 0.230 |
| Without NA12889 | 41 | 0.067 | 0.256 | 0.253 | 49 | 0.079 | 0.248 |
| Without NA12890 | 39 | 0.080 | 0.245 | 0.266 | 47 | 0.091 | 0.232 |
| Without NA12891 | 38 | 0.067 | 0.211 | 0.233 | 45 | 0.075 | 0.223 |
| Without NA12892 | 40 | 0.066 | 0.219 | 0.247 | 48 | 0.084 | 0.230 |
| Without trio A | 25 | 0.067 | 0.308 | 0.279 | 30 | 0.092 | 0.266 |
| Without trio B | 22 | 0.060 | 0.197 | 0.189 | 26 | 0.069 | 0.218 |

**Table S7:** The Y array. Calibration forms in five-fold cross-validation over the 59 training XY genomes, and comparison with Yruler on the 11 validation XY genomes and on all 70 XY genomes with assembled arrays (cenlen cross-validated on the training genomes). RMSE in kb; MedAPE, median absolute percentage error; single-core CPU times were measured on a matched compute host and include both counting and estimation for cenlen (Table S12).

|  | Training, 59 genomes |  | Validation, 11 genomes |  | All, 70 genomes |  |  |
|---|---|---|---|---|---|---|---|
| Method | RMSE | MedAPE (%) | RMSE | MedAPE (%) | RMSE | MedAPE (%) | CPU s per genome |
| *Calibration forms, cross-validated* |  |  |  |  |  |  |  |
| Zero-intercept linear | 49 | 2.51 | – | – | – | – | – |
| Linear with intercept | 46 | 3.82 | – | – | – | – | – |
| Quadratic | 48 | 2.96 | – | – | – | – | – |
| *Comparison with Yruler* |  |  |  |  |  |  |  |
| cenlen, Y-array estimate | 49 | 2.51 | 53 | 3.53 | 50 | 2.77 | 6.2 |
| Yruler, region mode | 52 | 2.52 | 46 | 2.31 | 51 | 2.49 | 50 |
| Yruler, whole file | – | – | – | – | – | – | 939 |

**Table S8:**
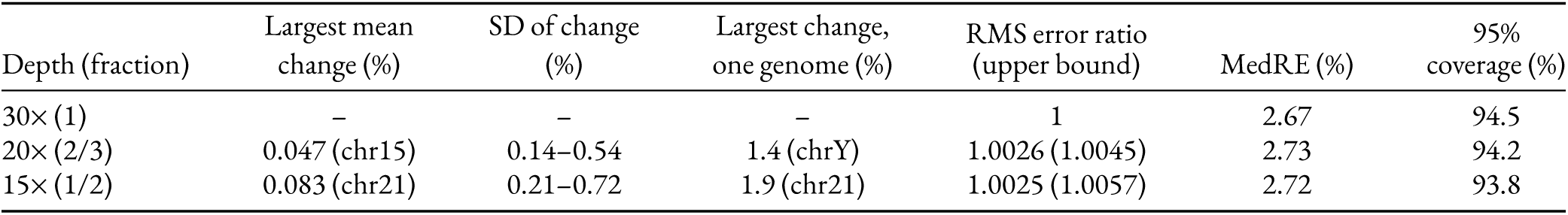
Coverage-downsampling sensitivity across 61 validation genomes. full depth (30×) and nested 2/3 (20×) and 1/2 (15×) subsamples: maximum mean relative shift across targets, range of per-target standard deviations, maximum single-genome shift, root-mean- square relative difference across all 566 genome–array pairs relative to full depth (with one-sided 95% upper bound), median absolute relative error and 95% prediction-interval coverage.

| Depth (fraction) | Largest mean change (%) | SD of change (%) | Largest change, one genome (%) | RMS error ratio (upper bound) | MedRE (%) | 95% coverage (%) |
|---|---|---|---|---|---|---|
| 30× (1) | – | – | – | 1 | 2.67 | 94.5 |
| 20× (2/3) | 0.047 (chr15) | 0.14–0.54 | 1.4 (chrY) | 1.0026 (1.0045) | 2.73 | 94.2 |
| 15× (1/2) | 0.083 (chr21) | 0.21–0.72 | 1.9 (chr21) | 1.0025 (1.0057) | 2.72 | 93.8 |

**Table S9:** Correlation of log array length between autosomes. Median Pearson *r* over pairs of the 22 autosomal arrays (231 pairs; 211 pairs with at least 15 genomes for the nested cross-validation rows). Within population, population means removed; relative, each array relative to the mean of the other 21 arrays within population, for which independent arrays of equal variance give about −1/21; trios, twice the covariance between a parent’s and the child’s arrays (2,000 trio bootstrap resamples for the interval).

| Estimate | Genomes | Median *r* |
|---|---|---|
| Assembled lengths | 279 nested cross-validation | 0.001 |
| cenlen out-of-fold estimates | 279 nested cross-validation | 0.014 |
| Out-of-fold log error | 279 nested cross-validation | 0.78 |
| cenlen estimates | 2,557 1000 Genomes | 0.057 |
| within population | 2,557 1000 Genomes | 0.032 |
| estimation covariance removed | 2,557 1000 Genomes | 0.008 |
| relative lengths | 2,557 1000 Genomes | −0.042 |
| Parent and child, different chromosomes | 427 trios | 0.003 (−0.009 to 0.017) |
| Parent and child, same chromosome | 427 trios | 0.946 |

**Table S10:** Target regions. The 3,432 intervals are released as a BED file on the reference of the 1000 Genomes CRAM files (GRCh38 with decoy and HLA sequences).

| Class | Definition |
|---|---|
| Centromere bands | merged UCSC cytoBand acen intervals for chr1–chr22 and chrX with 2 Mb flanks beyond adjacent heterochromatin; boundaries on chr1, chr2, chr6 and chrX extended distally by 0.7–3.0 Mb to prevent gap-boundary mismapping [11], merging five intervening gaps < 2 Mb |
| Heterochromatin, acrocentric short arms | intervals of the UCSC gap table |
| chrY | the whole contig (DYZ3 array and single-copy chrY k-mers); 0.87–1.61% of a CRAM file |
| Other contigs | 2,385 decoys; unlocalized, unplaced, alternative and HLA contigs; chrEBV |
| Unmapped reads | all |

**Table S11:** Technical validation of k-mer counting, array annotations and sample concordance.

| Comparison | Scope | Concordance |
|---|---|---|
| Streaming reference pipeline compared with meryl | 13,549,963 k-mers and total read, base and window counts across 78 CRAM files | exact concordance |
| cenlen whole-file pass compared with reference pipeline | all k-mer counts and summary totals across 85 CRAM files | exact concordance |
| Targeted container retrieval compared with whole-file pass | occurrences across chromosome-specific k-mer sets | ≥ 99.992% recovered |
| Replicate and network execution | duplicate local runs; local file compared with HTTP byte-range retrieval | exact concordance |
| Annotated versus sequence-extracted array span | 5,569 assembled haplotype arrays | exact agreement in all arrays |
| Short-read versus assembly sample matching | fraction of sample-identifying k-mers detected in matched reads versus three population-matched controls (thresholds: matched ≥ 0.80, margin ≥ 0.40) across 120 genomes | minimum matched fraction 0.966; minimum margin 0.461 |

**Table S12:** Per-genome runtime, memory and cloud HTTP byte-range benchmarks. (a) Per-genome CPU time, wall-clock time and peak resident memory across 30× CRAM benchmarks at three task scopes (CPU time sums across threads; CPU and wall-clock times report block means; peak memory reports the block maximum; CPU-time ratios compare unrounded baseline and cenlen means; ^a^streaming samtools fasta pipeline; ^b^wall-clock duration multiplied by CPU utilization in the workflow trace; ^c^benchmarked on the same compute host as HOROSCOPE, mean of two runs per file; ^d^matched Intel Xeon Gold 6430 replication hosts, with host B also executing the seven-genome reference-pipeline block; ^e^single run per file; ^f^eight-thread execution on the same host). (b) Targeted extraction on *All of Us* DRAGEN-aligned CRAM files in cloud storage (medians; 10th–90th percentiles in brackets).

| **a** Per-genome runtime and peak resident memory |  |  |  |  |
| Program | CPU time (s) | Wall time (s) | Peak memory (GB) | Speedup (×) |
|---|---|---|---|---|
| *All 25 targets, seven benchmark files* |  |  |  |  |
| Reference pipeline<sup>a</sup> | 1,324 | 407 | 0.3 | reference |
| cenlen count | 43.1 | 48.2 | 0.53 | 31 |
| cenlen estimate | 4.4 | 17.3 | 0.65 | – |
| cenlen count + estimate | 47.5 | 65.5 | 0.65 | 28 |
| *HOROSCOPE's 17 autosomes, six CEPH 1463 files* |  |  |  |  |
| HOROSCOPE k-mer extraction<sup>b</sup> | 11,060 | – | – | reference |
| cenlen, 25 targets<sup>c</sup> | 41.9 | 53.1 | 0.65 | 264 |
| cenlen, 17-autosome targets<sup>c</sup> | 35.3 | 43.8 | 0.65 | 313 |
| cenlen, 25 targets, host A<sup>d</sup> | 45.8 | 56.6 | 0.65 | 242 |
| cenlen, 17-autosome targets, host A<sup>d</sup> | 37.7 | 46.9 | 0.65 | 293 |
| cenlen, 25 targets, host B<sup>d</sup> | 49.1 | 68.0 | 0.65 | 225 |
| *Paired counts, seven benchmark files<sup>d</sup>* |  |  |  |  |
| cenlen count, released targets | 39.9 | 42.3 | 0.53 | – |
| cenlen count, 17-autosome targets | 33.7 | 35.6 | 0.53 | – |
| *Y-array benchmark, seven XY genomes<sup>c</sup>* |  |  |  |  |
| Yruler, region mode, one core | 49.6 | 57.0 | 0.74 | reference |
| Yruler, region mode, eight threads<sup>f</sup> | 69.3 | 18.1 | 1.65 | – |
| Yruler, whole file, one core<sup>e</sup> | 938.8 | 1000.7 | 2.39 | – |
| cenlen count, chrY contig | 3.7 | 3.9 | 0.53 | 13 |
| cenlen estimate from chrY counts | 2.5 | 7.5 | 0.65 | – |
| cenlen count + estimate, chrY | 6.2 | 11.5 | 0.65 | 8 |
| **b** Cloud HTTP byte-range retrieval on DRAGEN-aligned CRAM files |  |  |  |  |
| Quantity |  |  |  | Value |
| Fraction of targeted-extraction bytes on alternative contigs (DRAGEN vs 1000 Genomes) |  |  |  | 0.222 vs 0.025 |
| Fraction of targeted bytes saved by excluding alternative/HLA/EBV contigs |  |  |  | 0.21–0.23 |
| HTTP byte-range retrieval throughput (MB/s) |  |  |  | 34.9 [32.4–38.5] |
| HTTP byte-range retrieval wall-clock time per file (s) |  |  |  | 93.5 [82.3–113.6] |
| Ratio of ranged-HTTP to local-disk wall-clock time |  |  |  | 1.04 |
| Bytes transferred relative to whole-CRAM copy (DRAGEN-adapted list) |  |  |  | 12.6% |

**Table S13:** Ablation analysis of cenlen’s computational efficiency. CPU-time ratios between benchmark configurations on three high- coverage 30× CRAM files (Intel Xeon Gold 6430), from two experimental series (first two rows; component rows), each configuration executed twice in mirrored order. Each value is the CPU-time ratio between successive configurations; component rows ablate one engine feature during the targeted region pass. Conventional pipeline: samtools fasta piped into a k-mer counter with the same query table, over the whole file or over the target intervals with only the required fields decoded. CPU time is summed over threads; the counter ran on four threads in the first series and on one thread in the second, and the cenlen engine on one thread. The whole-file to engine ratio in the first series was 43.7–51.0 (count step only, shared node under concurrent load); the corresponding count-only ratio on a dedicated node across seven files was 31 (Table S12). All configurations produced identical k-mer counts.

| Optimization stage | HG00097 | HG01192 | NA18505 |
|---|---|---|---|
| Container and data-series selection (conventional tools) | 11.9 | 11.1 | 10.5 |
| Decode-coupled counting engine (same target regions) | 4.3 | 3.9 | 4.3 |
| *Engine components on the region pass (single-feature ablation)* |  |  |  |
| Decode-coupled counting in place of FASTA serialization and a separate k-mer counter | 2.1 | 2.0 | 1.9 |
| Engine slice reader and decoder with per-slice reference counting | 1.7 | 1.6 | 1.8 |
| 16-base Bloom prefiltering with per-slice reference counting | 1.5 | 1.5 | 1.5 |

**Table S14:** Shares of variance in 2,557 unrelated genomes of the 1000 Genomes Project on the haplotype scale. Shown with 95% bootstrap intervals: between superpopulations for log length and for log length relative to the genome’s mean over the other 21 autosomal arrays, and between populations within superpopulations for log length. *p*: 10,000 permutations of whole populations among superpopulations, for the log-length share (smallest attainable value 0.0001). SNV percentile: percentile of that share among 114,338 single common SNVs. Residual-bias check: superpopulation differences in mean out-of-fold training residuals within or exceeding half the inter-superpopulation median range; insufficient, fewer than two superpopulations with at least five training genomes. Genome mean: mean log length over the 22 autosomal arrays; common SNVs: ratio of summed components.

|  | Between superpopulations |  |  | Within | SNV | Residual-bias |
|---|---|---|---|---|---|---|
| Array | Log length | Relative to genome mean | *p* | superpopulation | percentile | check |
| chr1 | 0.010 (0.003–0.018) | 0.035 (0.024–0.046) | 0.0255 | 0.017 | 8 | exceeds |
| chr2 | 0.015 (0.008–0.024) | 0.010 (0.003–0.017) | 0.0049 | 0.013 | 13 | exceeds |
| chr3 | 0.005 (0.000–0.012) | 0.023 (0.014–0.034) | 0.0507 | 0.010 | 3 | within |
| chr4 | 0.047 (0.034–0.061) | 0.010 (0.003–0.018) | 0.0001 | 0.011 | 48 | within |
| chr5 | 0.030 (0.019–0.042) | 0.040 (0.028–0.053) | 0.0012 | 0.017 | 29 | exceeds |
| chr6 | 0.010 (0.003–0.018) | 0.003 (-0.002–0.009) | 0.0051 | 0.014 | 8 | within |
| chr7 | 0.091 (0.073–0.111) | 0.090 (0.072–0.110) | 0.0001 | 0.012 | 79 | within |
| chr8 | 0.012 (0.005–0.020) | 0.019 (0.011–0.027) | 0.0075 | 0.010 | 9 | exceeds |
| chr9 | 0.046 (0.034–0.058) | 0.028 (0.018–0.040) | 0.0001 | 0.011 | 47 | within |
| chr10 | 0.077 (0.060–0.094) | 0.054 (0.039–0.069) | 0.0001 | 0.013 | 71 | within |
| chr11 | 0.068 (0.052–0.084) | 0.022 (0.013–0.032) | 0.0001 | 0.005 | 65 | within |
| chr12 | 0.017 (0.009–0.027) | 0.027 (0.016–0.038) | 0.0011 | 0.010 | 15 | exceeds |
| chr13 | 0.046 (0.034–0.060) | 0.047 (0.035–0.060) | 0.0001 | 0.010 | 48 | exceeds |
| chr14 | 0.037 (0.025–0.049) | 0.083 (0.066–0.099) | 0.0001 | 0.005 | 38 | within |
| chr15 | 0.039 (0.027–0.052) | 0.025 (0.015–0.037) | 0.0001 | 0.017 | 40 | within |
| chr16 | 0.086 (0.070–0.103) | 0.142 (0.123–0.163) | 0.0001 | 0.026 | 76 | within |
| chr17 | 0.089 (0.074–0.106) | 0.036 (0.026–0.048) | 0.0001 | 0.006 | 78 | exceeds |
| chr18 | 0.015 (0.008–0.024) | 0.010 (0.004–0.017) | 0.0059 | 0.014 | 13 | insufficient |
| chr19 | 0.044 (0.032–0.058) | 0.010 (0.003–0.018) | 0.0001 | 0.010 | 46 | exceeds |
| chr20 | 0.087 (0.070–0.105) | 0.051 (0.038–0.066) | 0.0001 | 0.008 | 77 | within |
| chr21 | 0.171 (0.147–0.196) | 0.168 (0.146–0.191) | 0.0001 | 0.005 | 96 | within |
| chr22 | 0.045 (0.033–0.057) | 0.012 (0.005–0.020) | 0.0001 | 0.011 | 46 | within |
| chr1+chr5+chr19 | 0.041 (0.027–0.054) | – | 0.0010 | 0.030 | 42 | insufficient |
| chr13+chr21 | 0.209 (0.189–0.231) | – | 0.0001 | 0.012 | 98 | within |
| chr14+chr22 | 0.022 (0.013–0.031) | – | 0.0006 | 0.011 | 20 | exceeds |
| chr1+chr5+chr16+chr19 | 0.046 (0.034–0.060) | – | 0.0009 | 0.037 | 48 | insufficient |
| Genome mean | 0.157 (0.138–0.178) | – | 0.0001 | 0.066 | 95 | exceeds |
| Common SNVs | 0.0682 | – | – | 0.007 | – | – |

**Table S15:** Genome-mean log length by superpopulation. Difference from the overall mean (%): nested cross-validation genomes with at least 12 assembled autosomes, and the 2,557 unrelated 1000 Genomes genomes.

|  |  | Assembled genomes |  |  | 1000 Genomes |
|---|---|---|---|---|---|
| Superpopulation | *n* | Assembled | Estimated | Residual | Estimated |
| AFR | 25 | −1.0 | +0.7 | +1.8 | −1.8 |
| AMR | 9 | −4.8 | −5.8 | −0.9 | −3.3 |
| EAS | 8 | +5.3 | +1.0 | −4.2 | +2.2 |
| EUR | 7 | −0.3 | +0.3 | +0.7 | −2.9 |
| SAS | 6 | +4.8 | +4.2 | −0.6 | +5.6 |

**Table S16:** Association between X-array length and *MED14* alternative splicing. Lead intron chrX:40,666,851–40,668,597, cluster clu_50837, ∼20 Mb from the X centromere; 730 MAGE donors, 388 X-linked genes tested. Local-variant adjustment conditions on the three SNVs within 1 Mb of *MED14*’s transcription start site most strongly associated with intron excision (forward selection across 3,763 common variants with minor allele frequency ≥ 0.05).

| Model | Donors | *r* | *p* | Note |
|---|---|---|---|---|
| All donors | 730 | 0.168 | 7.1 × 10<sup>−6</sup> | gene-level *p* = 8.1 × 10<sup>−5</sup>, *q* = 0.028; sole X-linked gene at *q* < 0.05 in the whole-chromosome scan |
| Females | 381 | 0.223 | 2.0 × 10<sup>−5</sup> |  |
| Males | 349 | 0.103 | 0.061 | sex difference *p* = 0.10 (Fisher *z*) |
| Adjusted for genome mean array length | 730 | 0.164 | 1.1 × 10<sup>−5</sup> | females 0.218, males 0.100 |
| Adjusted for three local variants | 730 | 0.157 | 2.65 × 10<sup>−5</sup> | 94% of unadjusted *r* |

